# Structure, Dynamics and Free Energy Studies on the Effect of Spot Mutations on SARS-CoV-2 Spike Protein Binding with ACE2 Receptor

**DOI:** 10.1101/2023.07.19.549772

**Authors:** George Rucker, Hong Qin, Liqun Zhang

**Affiliations:** Chemical Engineering Department, Tennessee Technological University, Cookeville, TN, 38501; Computer Science department, University of Tennessee Chattanooga, Chattanooga, TN 37403; Chemical Engineering Department, University of Rhode Island, Kingston, RI, 02881

## Abstract

The ongoing COVID-19 pandemic continues to infect people worldwide, and the virus continues to evolve in significant ways which can pose challenges to the efficiency of available vaccines and therapeutic drugs and cause future pandemic. Therefore, it is important to investigate the binding and interaction of ACE2 with different RBD variants. A comparative study using all-atom MD simulations was conducted on ACE2 binding with 8 different RBD variants, including N501Y, E484K, P479S, T478I, S477N, N439K, K417N and N501Y-E484K-K417N on RBD. Based on the RMSD, RMSF, and DSSP results, the overall binding of RBD variants with ACE2 is stable, and the secondary structures of RBD and ACE2 are consistent after the spot mutation. Besides that, a similar buried surface area, a consistent binding interface and a similar amount of hydrogen bonds formed between RBD with ACE2 although the exact residue pairs on the binding interface were modified. The change of binding free energy from spot mutation was predicted using the free energy perturbation (FEP) method. It is found that N501Y, N439K, and K417N can strengthen the binding of RBD with ACE2, while E484K and P479S weaken the binding, and S477N and T478I have negligible effect on the binding. Spot mutations modified the dynamic correlation of residues in RBD based on the dihedral angle covariance matrix calculation. Doing dynamic network analysis, a common intrinsic network community extending from the tail of RBD to central, then to the binding interface region was found, which could communicate the dynamics in the binding interface region to the tail thus to the other sections of S protein. The result can supply unique methodology and molecular insight on studying the molecular structure and dynamics of possible future pandemics and design novel drugs.

## Introduction

COVID-19 pandemic which is the result of infection by SARS-Coronavirus-2 (CoV-2), had claimed over 6 million lives as of June 2023, and continues impacting worldwide health [1]. CoV- 2 expresses S (Spike) protein [2,3], which is responsible for binding to the host cell receptor followed by fusion of the viral and cellular membranes, respectively [4]. To engage a host cell receptor, the receptor-binding domain (RBD) of the S protein undergoes hinge-like conformational movements [5], thus mediates interaction with the receptor angiotensin-converting enzyme 2 (ACE2) [6,7,8,9] (the RBD and ACE2 binding structure is shown in Figure 1 (a)). The binding of RBD with ACE2 is believed to be the critical initial event in the infection cascade. The interface residues on RBD thus play key roles in the spike protein’s binding with ACE2.

**Figure 1.**
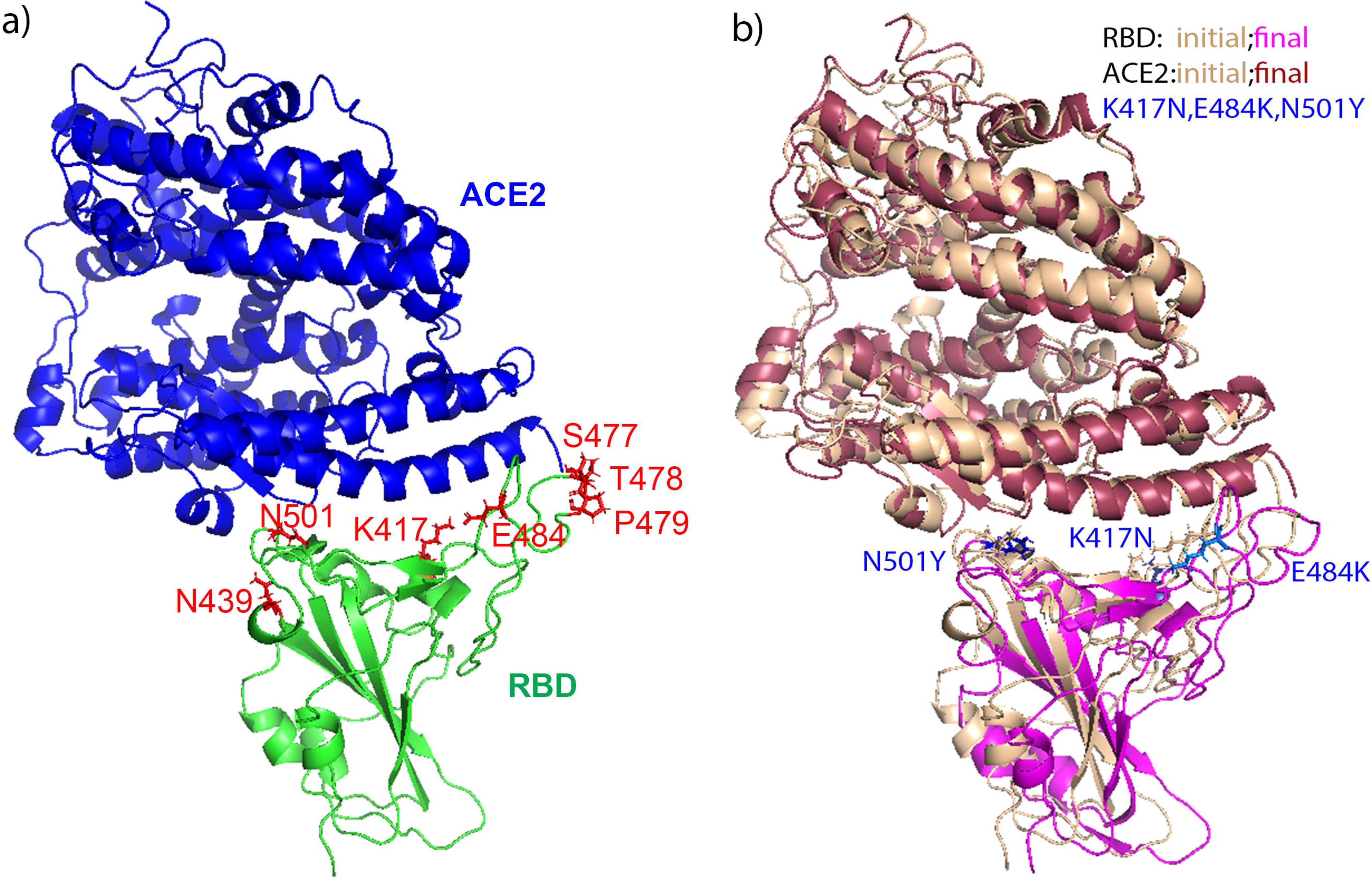
The binding structure of ACE2 (in blue) with RBD (in green) (a), with the mutations (shown in red sticks) on RBD labeled; and the comparison of the initial and final structures of ACE2 bound with RBD-trimutant variant (b). In b), the initial structures of ACE2 and RBD are shown in wheat, and the final structure of ACE2 is shown in raspberry and magenta for RBD. The mutations of E484K, K417N, and N501Y are shown in blue sticks in both the initial and final structures.

Coronaviruses are enveloped viruses that possess a positive-sense single-stranded RNA genome 26–32 kb in length [10], and have a remarkable mutation rate that allows them to evolve in a way that perhaps by increasing their transmissibility [11,12, 13, 14] and/or their resistance to protective immunity induced by previous infection or vaccines [15, 16, 17, 18, 19, 20]. Up to now, different variants of concern (VOCs) have shown up, including Alpha, Beta, Gamma, Delta, and Omicron [21,22,23,24,25,26,27,28] (Some mutations involved are shown in Table S1 in SI). The Alpha variant was found in UK in September of 2020, while Beta variant was found in North Africa in May of 2020. Gamma variant was found in Brazil [21,22,23,25]. The Omicron variant which contains 11 mutants on RBD, is still the dominant cause of the resurgence of infection in many countries currently [27]. Most of the mutations (around 58% of mutations) on RBD are located at the receptor binding motif which is the binding interface with ACE2 [29].

N501Y on RBD, which appeared in Alpha, Beta, Gamma and Omicron variants, increased the binding affinity of RBD with ACE2 [30, 31] and enhanced the viral resistance to neutralizing antibodies [32]. S477N appeared in Omicron and E484K appeared in both Beta and Gamma variants, were reported to enhance the binding affinity of S protein with ACE2 [33,34]. K417N found in both Beta and Omicron variants was reported to decrease the binding affinity of RBD and ACE2 [35], and was thought to cause immune escape [36]. The triple mutant of N501Y-E484K- K417N appeared in Beta variant, has been shown to increase the infectivity of virus considering the effect of each single mutant [35]. N439K, which was found in multiple regions in the world and a quite common mutation, can enhance the binding of RBD with human ACE2 receptor while evade antibody mediated immunity [37]. P479S, which also happened with high frequency at most countries based on SARS-CoV-2 genome samples deposited at GISAID in early 2020 [38], was predicted to make the virus less infective based on algebraic topology-based machine learning modelling [29]. Besides that, based on the report of most frequently occurring mutations in RBD in January 2021 (https://www.gisaid.org/hcov19-mutation-dashboard), T478I which appeared in Alpha variant, however did not influence the binding of RBD and ACE2 [39]. Those mutants all appeared on the RBD binding interface.

Besides those experimental work, in order to find out the mechanism of CoV-2 variants binding with ACE2 and to design effective therapeutics to deal with CoV-2 variants, molecular dynamics simulation methods have been applied to study RBD binding with ACE2 [33,40, 41,34, 42, 43,44,45,46,47]. Different simulation methods were used to predict the binding free energy change (ΔΔG) [48, 49], including free energy perturbation (FEP) method [43,41], molecular mechanics Poisson-Boltzmann surface area (MM-PBSA) method [42], molecular mechanics-generalized Born surface area (MM/GBSA) method [45] and so on. Li et al. [49] and other researchers found that FEP method can predict ΔΔG reasonably well [41,48, 50] and better than other methods [49]. Thus FEP method was applied to predict the spot mutation on RBD to the binding of RBD variants with ACE2 in this work. Besides that, dynamic correlation was calculated to predict correlation matrix and dynamic network in RBD in order to understand how the S spike mutations on RBD binding interface can affect the binding of RBD with ACE2. Since spot mutations happened quite often on CoV-2 RBM [29], a comparative study on multiple single mutations on RBD binding with ACE2 is necessary. In total 7 RBD single-site mutations which happened naturally and can potentially affect the binding and dynamics of RBD with ACE2 were studied including N501Y, E484K, K417N, N439K, P479S, T478I, S477N, and one triple mutant N501Y-E484K-K417N.

Their effects on the binding of RBD with ACE2 were analyzed. Besides all-atom NAMD simulations, FEP method was applied to predict ΔΔG of RBD with ACE2 coming from the spot mutation. The simulation predictions helped to clarify the contradicting available experimental results. Calculating the dihedral angle covariance matrices and doing dynamic network analysis, an intrinsic network community was predicted. Our results revealed the molecular mechanism that underlies the dynamics and energy change of some RBD variants binding with ACE2. Such kind of information will be valuable for the development of further vaccines and neutralizing antibodies against mutant forms of the SARS-CoV-2 virus.

## Materials and Methods

1). Based on the survey up to now, 7 naturally happened single RBD mutants were studied: E484K, N439K, P479S, T478I, N501Y, K417N, S477N, besides one RBD triple mutant which has E484K, N501Y and K417N. The selection of those mutations is based on different SARS-CoV-2 variants shown as listed in Table S1 in SI. The mutation sites on RBD and ACE2 binding interface are shown in Figure 1(a). The crystal structure of the RBD domain of the Spike protein is available in complex with ACE2 at 2.45 Å resolution in the PDB with ID 6M0J [51], which was used as the starting structure to build the RBD mutants bound with ACE2. VMD program [52] was applied to mutate residues on RBD to generate RBD mutants. The corresponding RBD and ACE2 structures in [51] were used in reference simulations of RBD and ACE2 in free forms respectively. The bound and free forms of simulations conducted in this project are listed in Table S2.

Simulation systems on RBD mutants bound with ACE2 were set up using VMD program and CHARMM36m forcefield [53]. Each system was solvated with enough amount of TIP3P water molecules with the minimum distance between protein and water box edge being at least 12 Å, and was neutralized with 0.15M NaCl solvent. After a brief energy minimization using the conjugate gradient and line search algorithm, 4 ps of dynamics was run at 50 K, and then the system was brought up to 310 K over an equilibration period of 1 ns running all-atom molecular dynamics simulations using NAMD program [54] and CHARMM36m forcefield [53]. The temperature was 310K and a desired pressure of 1 atm using standard thermo-and barostats. The sampling run was 100 ns with a time step of 1 fs for each system. The trajectory output frequency was 1 ps. To do comparisons with the binding of RBD mutants with ACE2, the ACE2 bound with RBD wildtype system, the ACE2 only in solvent system, and the RBD only in solvent system were also set up and 100 ns simulation was also performed for them each. The simulation systems, set up, the number of atoms and box size information are shown in Table S2.

To analyze the trajectories, the Root Mean Square Deviation (RMSD) and Fluctuations (RMSF) of the proteins were calculated using the VMD program and an in-house analysis script based on the coordinates of the backbone CA atoms after aligning the trajectories respectively, to the original crystal structure of the RBD, ACE2, and to the initial complex structure of the RBD and ACE2 from Lan et al.[51]. The buried surface area (BSA) for the complex was calculated in two steps using the VMD program and a script using the Richards and Lee method with the water probe size of 1.4 Å (Lee and Richards, 1971). First, the total solvent accessible surface area of the complex (ASA_COMPLEX_) was calculated based on the complex’s trajectory. Second, the accessible surface area of each protein in the complex (ASA_RBD_, ASA_ACE2_) was calculated for each protein individually. Then, the buried surface area (BSA) is calculated using Equation (1):

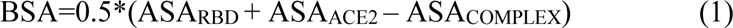

The number of hydrogen bonds between the RBD and ACE2 were calculated using the VMD program with the heavy atom distance cutoff of 3.0 Å and the angle cutoff of 20 degrees deviation from H-bond linearity. The time a particular H-bond is formed over the course of the simulation is monitored and is expressed as % occupancy. In order to find out the residues on the binding interface, the closest distance between every residue atom (including hydrogen) between the RBD and ACE2 was calculated and averaged over the trajectory run. The average distances between each residue on RBD and on ACE2 are shaded by proximity on a red to white color-scale and were used to build the distance maps.

2). Dihedral Angle Correlation, network community calculation. To quantitatively describe the motional correlations of the backbone in RBD, we calculated the ϕ and ψ cross-correlation matrix for all the residues except the first or last one on the sequence. ϕ describes the rotation of the N−CA bond and involves the CO−N−CA−CO bonds. ψ describes the rotation of CA and the CO bond and involves the N−CA−CO−N bonds. Both ϕ and ψ were calculated for residues at different times using the wordom program [55,56]. Because both ϕ and ψ are angular variables, to avoid the periodicity problem, the circular correlation coefficient, which is a T-linear dependence, was calculated in this project following the method of Fisher [57,58] and Mardia and Jupp[59], and in the same way as in our previous work [60,61]. The ϕ−φ cross-correlation coefficients (matrices) were calculated of all combinations of dihedral angles. In order to simplify the data representation, the ϕ−ϕ, ϕ−φ, and φ−φ correlation matrices were combined and the element with the largest magnitude at residue pair position was always chosen to build the final matrix.

In the dynamic network community calculation, the network model was built by the NetworkView plug-in of VMD [62] and the program Carma [63]. In the network, the nodes could be considered as a single atom or a cluster of atoms. In this project, each CA atom in one amino acid was treated as one node. The edge between two nodes was defined with the cutoff distance of 4.5Å for at least 75% of molecular dynamics simulated trajectory.

3). In order to find out the contribution of mutant residues on RBD to its binding with ACE2, FEP method was applied in the same way as in [50]. The change of free energy of each mutation to the binding was predicted, including E484K, N439K, N501Y, P479S, S477N, T478I, K417N on RBD binding with ACE2.

In total, two kinds of systems were set up. One is the RBD in wildtype form in solvent (unbound state). The other is the RBD mutant bound with ACE2 (bound state). For both kinds of systems, mutation on one residue (E484K, N439K, K417N, T478I, P479S, S477N, N501Y) on RBD (mutant systems) was conducted using VMD program [52]. After solvating the unbound and bound systems with RBD in both wildtype and mutant forms with enough amount of water and neutralizing them with 0.15M NaCl, NAMD all-atom molecular dynamics simulations were performed to equilibrate both kinds of systems using an NVT ensemble after a brief energy minimization. The dual-topology methodology [64,65,66], and the soft-core potentials [67,68] were applied to overcome endpoint singularities [69] in the FEP simulation, following the same method as in our previous work [50]. The free energy change in mutation for residue *i* in thermodynamic cycle can be calculated using Equation (2):

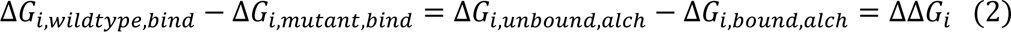

Here, *ΔΔG_i_* is the relative change of alchemical free energy of residue *i* during the binding of RBD with ACE2 as the thermodynamic cycle shown in Figure S1. In each FEP simulation, 3 ns simulations were performed in each window for both forward and backward directions and for both the bound and free states after 0.5 ns simulations were conducted to equilibrate the system. In this project, the interaction calculation includes 10 windows, ranging from a thermodynamic coupling parameter λ values of zero to 1 (with a gap of 0.1) for a total of 20 simulations per mutation. The time step was 1.0 fs.

## Results

### 1). RMSD, RMSF, ΔRMSF, DSSP results for ACE2 and RBD variants

#### RMSD result

Based on all-atom NAMD simulations (100 ns each) on different systems, the RMSDs of the complex, RBD, ACE2 were calculated after aligning the trajectories to the RBD wildtype bound with ACE2, RBD wildtype, and ACE2 individually. RBD mutants, ACE2, and the complexes are very stable during 100 ns simulations and are very consistent with those from ACE2 bound with RBD wildtype (as the RMSD vs simulation time result shown in Figure S2 in SI). Calculating the average RMSD of the complex, the RBD, and the ACE2, the results are shown in Table 1. Only ACE2&RBD-N501Y complex has a RMSD (2.09±0.2Å) lower than that of the ACE2&RBD wildtype(2.21±0.4Å). Although RBD-N501Y has the highest RMSD (1.54±0.1Å), ACE2 bound with this RBD variant has the lowest RMSD (1.84±0.2Å). That suggests that RBD- N501Y can fit the binding pocket of ACE2 with the minimum structure change on ACE2 and the maximum structure adaptation of RBD-N501Y. On the other hand, the complexes of ACE2 with RBD-trimutant, RBD-T478I, and RBD-N439K showed the largest structural deviation from the wildtype(3.05±0.4Å, 3.06±0.4Å, 3.34±0.5Å). When bound with RBD-N439K, ACE2 has the largest structural deviation (2.60±0.3Å) comparing to the ACE2 wildtype (1.88±0.2Å). The binding of RBD with ACE2 decreased the structural deviation of both RBD and ACE2. Overall, the RMSD result suggests that spot mutation did not affect the binding structure of ACE2 with RBD significantly.

**Table 1.** The average and standard deviations of RMSDs of ACE2&RBD complex, RBD wildtype and variants, and ACE2, the number of hydrogen bonds (hbonds) formed, and the buried surface area (BSA) between ACE2 and RBD. The standard deviations were calculated based on the second-half of NAMD simulations.

| Simulations | RMSD<sub>Complex</sub><br>(Å) | RMSD<sub>RBD</sub><br>(Å) | RMSD<sub>ACE2</sub><br>(Å) | Hbond | BSA<br>(Å<sup>2</sup>) |
| --- | --- | --- | --- | --- | --- |
| E484K&ACE2 | 2.40±0.3 | 1.22±0.1 | 1.92±0.1 | 2.63±1.7 | 864.3±57.4 |
| N439K&ACE2 | 3.34±0.5 | 1.18±0.1 | 2.60±0.3 | 2.69±1.4 | 876.0±46.8 |
| N501Y&ACE2 | 2.09±0.2 | 1.54±0.1 | 1.84±0.2 | 2.62±1.3 | 843.6±54.4 |
| P479S&ACE2 | 2.74±0.7 | 1.21±0.1 | 2.11±0.2 | 3.23±1.5 | 861.0±62.0 |
| K417N&ACE2 | 2.68±0.3 | 1.20±0.1 | 2.45±0.3 | 2.32±1.4 | 838.1±45.5 |
| S477N&ACE2 | 2.89±0.4 | 1.15±0.1 | 2.50±0.2 | 3.03±1.4 | 908.6±41.4 |
| T478I&ACE2 | 3.06±0.4 | 1.22±0.1 | 2.43±0.2 | 1.49±1.1 | 729.7±50.3 |
| Tri-mutant&ACE2 | 3.05±0.4 | 1.32±0.2 | 2.10±0.2 | 2.01±1.2 | 835.5±46.5 |
| RBD&ACE2 wt | 2.21±0.4 | 1.23±0.1 | 1.88±0.2 | 2.64±1.4 | 863.3±44.4 |
| RBD wt, free |  | 1.52±0.2 |  |  |  |
| ACE2 free |  |  | 2.62±0.4 |  |  |

#### RMSF and ΔRMSF result

Calculating the Root Mean Squared Fluctuation (RMSF) of RBD and ACE2 from different simulations, the results are shown in Figure 2(Left) and (Right).

**Figure 2.**
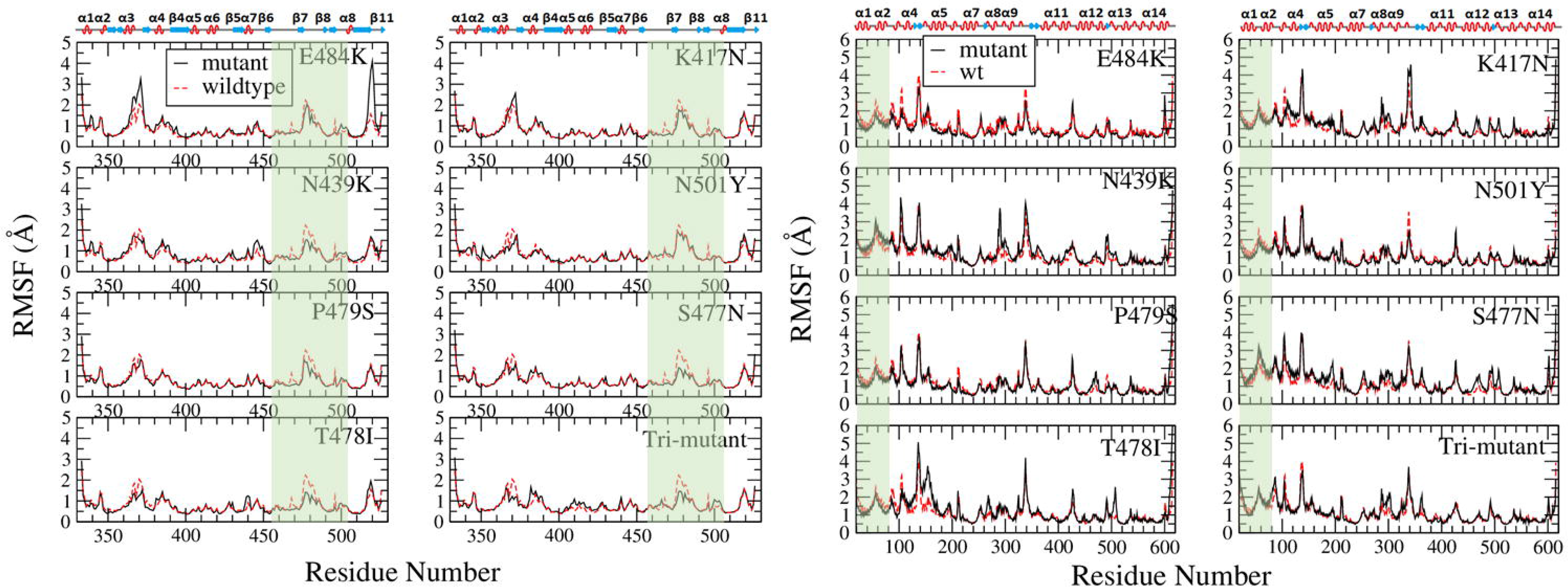
Comparison of RMSF from RBD variants and wildtype (Left) and ACE2 (Right). The variant results are shown in black solid line while the wildtype shown in red dashed line. The binding regions on RBD and ACE2 are shaded in light green, and the secondary structures of RBD and ACE2 are shown on the top of RMSF figures.

Residues from 455 to 505 on RBD bound with ACE2 at the region of RES19 to RES81, which are highlighted in light green for RBD on Figure 2(Left) and for ACE2 on Figure 2(Right).

Within the binding region, all the mutants have a smaller RMSF than the wildtype, especially S477N, N439K, T478I and the tri-mutant. However, at other regions such as RES360 to RES390, E484K showed a larger structure fluctuation.

Mapping the change of RMSF (ΔRMSF=RMSF_mutant_-RMSF_wt_) on the secondary structure of RBD and ACE2, the result for N501Y is shown in Figure 3(Left) and the tri-mutant bound with ACE2 in Figure 3(Right). Comparing to the wildtype, RBD-N501Y is slightly more rigid at RES501 site and RES484 site, although other binding region has a similar rigidity to the wildtype form. However, ACE2 is more rigid on the binding interface except the loop region which becomes more flexible than the ACE2 when bound with RBD wildtype. Figure 3(Right) shows that part of the binding interface on RBD becomes more rigid, part unchanged while the left region become more flexible comparing to the RBD wildtype, while ACE2 also has part of the interface more rigid, part unchanged, and other part more flexible comparing to the ACE2 bound with RBD wildtype. Those are consistent with RMSF result in Figure 2. Similarly, the ΔRMSF of other RBD mutants and ACE2 were calculated and mapped to the secondary structure of RBD and ACE2 (shown in Figure S3 to S5). In summary, the binding interface of ACE2 molecule becomes more rigid when bound with RBD variants except with S477N and N439K which have more flexible region than rigid region on the binding interface. Comparing to the RBD wildtype, RBD variants only become slightly more rigid except RBD-N439K variant which becomes mostly more flexible. Faraway regions of ACE2 bound with RBD variants usually become more flexible (such as RBD-S477N, RBD-N439K, and RBD-K417N, RBD-T478I), but not for the ACE2 bound with E484K, which overall become more rigid. Those can agree with RMSF results shown in Figure 2, and the result suggests that spot mutations on RBD can change the overall structure flexibility of RBD and ACE2 comparing to the wildtype. Such kind of change of flexibility should contribute to the adaptation of RBD variants to fit the binding pocket of ACE2 which also can adjust its flexibility accordingly.

**Figure 3.**
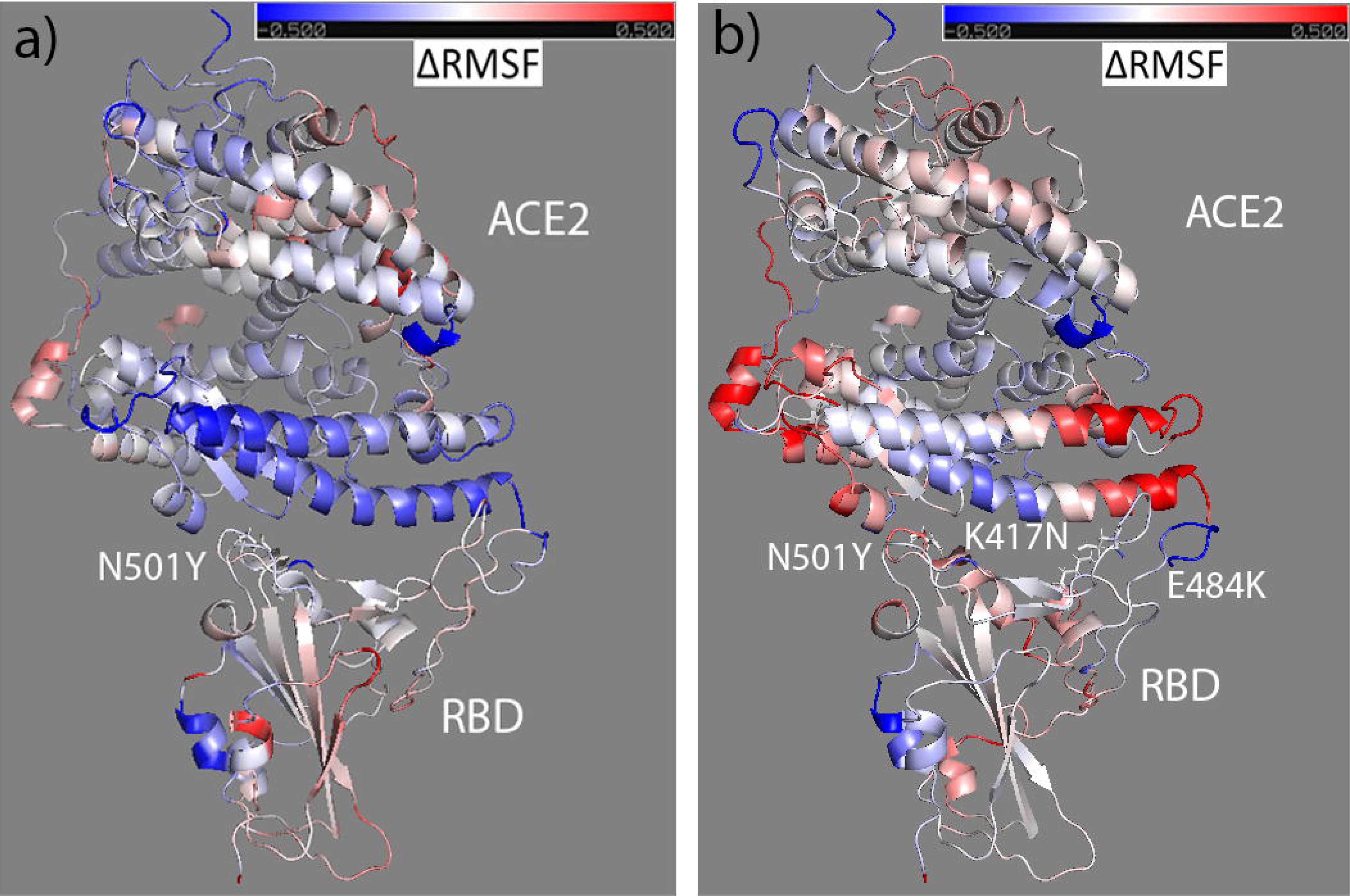
The ΔRMSF of RBD-N501Y and ACE2 (Left) from RBD wildtype to variant forms mapped to the initial bound structure of ACE2&RBD-N501Y; and the ΔRMSF of RBD- trimutant and ACE2 (Right) mapped to its initial bound structure of the complex. Colors range from blue to white to red (with white representing no change in RMSF, blue representing an RMSF decrease, and red representing an RMSF increase). The ΔRMSF is in the range from -0.5 (totally blue) to 0.5 (totally red).

#### DSSP result

In order to track the structure change of ACE2 and RBD contributed by the spot mutation, the evolution of the secondary structure was predicted based on DSSP [70] calculation over the simulation time using the wordom program. The DSSP program assigns the protein secondary structure to one of the nine states defined, including three helix types (3_10_, G; α, H; π, I), two extended sheet types (antiparallel and parallel, E and X), two types of turns including the helix turn (T) and β bridges (B), bend (S), and others (L). The secondary structure was calculated for each residue at each time frame during the simulation time. Based on that, the average proportion of helix (including G, H and I), sheet (including E), turn (including B and T) and other (including X, S and L) structures on RBD and ACE2 over the simulation time were calculated, and the results in comparison with available literature data [39] are shown in Figure 4 for RBD and in Figure S6 in SI for ACE2.

**Figure 4.**
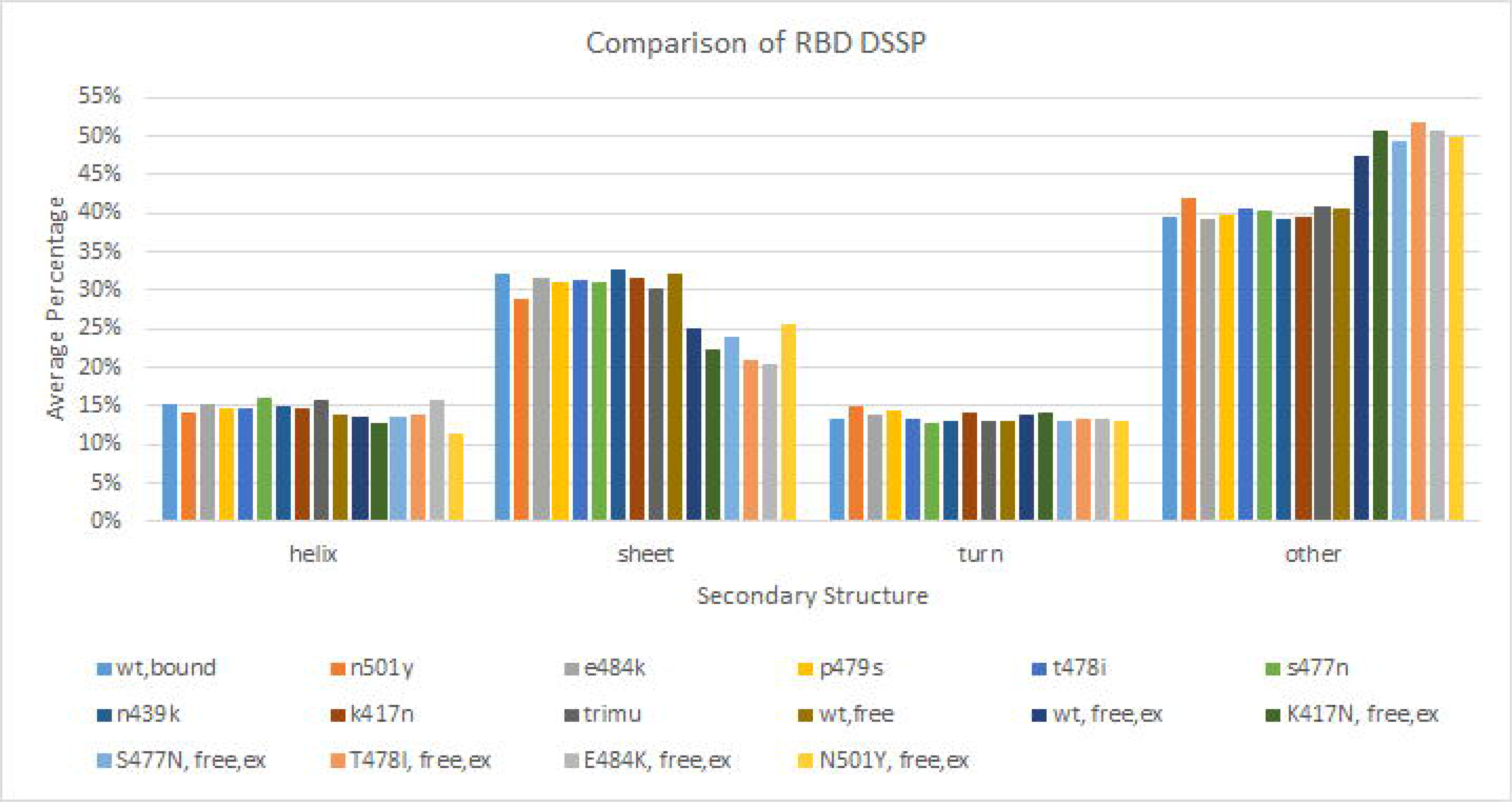
Comparison of RBD DSSP with available literature data.

Overall, the proportions of RBD secondary structure elements are very consistent with available experimental data [39], with around 13%-15% of helix, 30%-32% of sheet (including both parallel and anti-parallel), and 13%-14% of turns, and 40%-43% of others. That implies the RBD structures during NAMD simulations are reasonable. In Figure 4, comparing the DSSP of RBD mutants with RBD wildtype from the experimental data (the last 6 columns which only have RBD in free state available), it shows that the average percentage of other structure is higher in RBD mutants than the RBD wildtype. That is consistent with the simulation prediction for RBD wildtype and variants (the first 9 columns which are all in bound form). That suggests that spot mutation can increase undefined structure (other structure) in RBD. Although DSSP of RBD in wildtype free state (wt, free) has similar helix and turn ratios to the experimental data of RBD in wildtype free state (wt, free, ex), its sheet ratio is higher than the experimental data by around 6% while its other structure ratio is lower than the experimental data by around 6%. Such kind of discrepancy should come from the different definitions of sheet from Anderson [70] and the experimental method. Because of that, the average percentage of sheet structure from simulations are overall higher than experimental data while the average percentage of other structures are overall lower than experimental data.

Comparing results from simulation only or from experiments only, overall spot mutation only changes the secondary structure distribution of RBD slightly in both bound and free states. It almost does not change the secondary structure distribution of ACE2 as shown in Figure S6. So, overall, the mutants and wildtype have similar secondary structure distribution and spot mutation should only affect the secondary structure locally.

### 2). Hydrogen bonds formed on the binding interface

Forming intermolecular hydrogen bonds is one of the major driving forces for the binding of RBD with ACE2. Calculating the number of hydrogen bonds formed between ACE2 and RBD mutants over the simulation time (shown in Figure S7), overall RBD mutants formed similar amount of hydrogen bonds with ACE2 to the RBD wildtype during the simulation time. Calculating the average and standard deviation of the number of hydrogen bonds formed, the results are shown in Table 1. RBD can form averagely 1.5 to 3.2 hydrogen bonds with ACE2. S477N and P479S formed more hydrogen bonds with ACE2 than the wildtype, while T478I formed the least, and the tri-mutant formed the second to the least amount of hydrogen bonds with ACE2. Tracking the residue pairs on the binding interface, the results are shown in Table S3 in SI. There are 8 pairs of residues forming hydrogen bonds on the RBD wildtype-ACE2 interface, including Gly502-Lys353, Tyr505-Glu37, Asn487-Tyr83, Thr500-Tyr41, Lys417-Asp30, Gln493-Glu35, Tyr449-Asp38, Gln498-Gln42. With the triple-mutation, three pairs (Lys417-Asp30, Tyr449-Asp38, Gln498- Gln42) were broken and only the other five still formed hydrogen bonds on the tri-mutant&ACE2 interface. However, a new hydrogen bond formed between THR500-ASP355, and the interface structures generated from ligplot program [71] in comparison with RBD wildtype are shown in Figure 5. The details of hydrogen bonds formed between ACE2 and RBD wildtype and variants are shown in Table S3. Overall, similar binding interface formed in RBD wt&ACE2 and RBD variant&ACE2, although the exact residues forming hydrogen bonds changed and the number of hydrogen bonds formed also changed by certain amount. Such kind of small modification on the binding interface contributes to the different binding free energy of ACE2 and RBD which mainly comes from hydrogen bonds formation and electrostatic and hydrophobic interactions [45], and affects the infectivity of CoV-2 variants.

**Figure 5.**
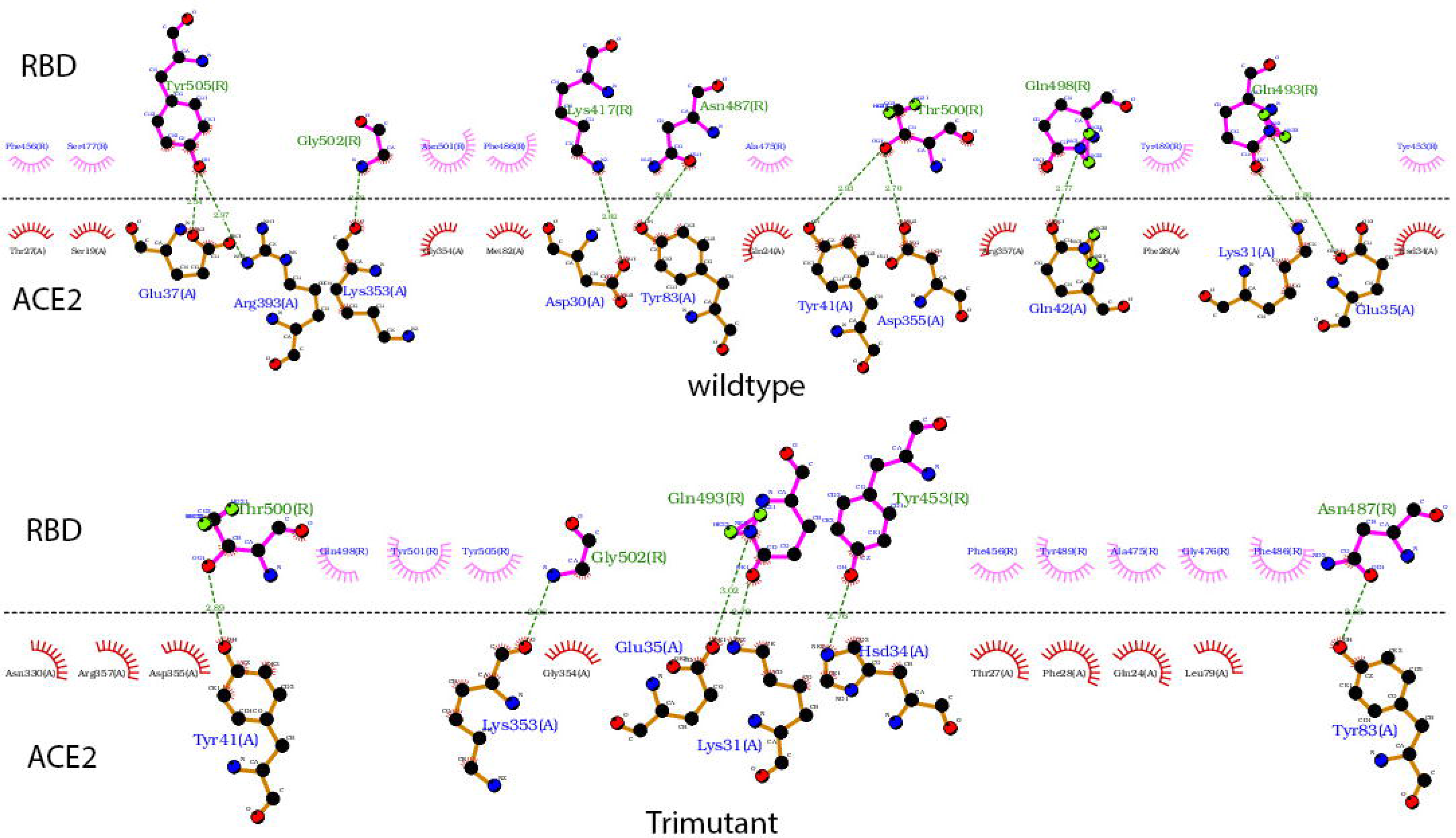
Hydrogen bonds formed on the ACE2&RBD interface for the wildtype (Top) and tri-mutant (Bottom) generated from ligplot program. Residues formed hydrogen bonds are connected using dashed green lines. Residues on RBD are labelled in light green with bonds shown in magenta, while residues on ACE2 are labelled in blue with bonds shown in orange.

### 3). BSA and distance map result

#### BSA comparison

Calculating the buried surface area (BSA) between ACE2 and RBD variants over the simulation time, the results in comparison with the RBD wildtype are shown in Figure 6. RBD wildtype and variants can bind with ACE2 consistently during the simulation time. Calculating the average and standard deviations of BSA based on the second half of the simulation time, the results are shown in Table 1. Overall, the RBD variants and ACE2 form a similar BSA as the RBD wildtype and ACE2. The RBD-T478I formed a slightly smaller BSA than other mutants, while RBD-E484K, RBD-N439K and RBD-S477N form a larger BSA with ACE2 than the RBD wildtype; the RBD-trimutant forms the smallest BSA among all the simulations especially during 20 to 100 ns period in the simulations as shown in Figure 6. Based on the findings from Chan et al. [43], a larger BSA suggests a stronger binding of RBD with ACE2, thus it is expected that E484K, N439K and S477N variants should form a stronger binding with ACE2 than other variants, while the tri-mutant should form the less stable binding with ACE2.

**Figure 6.**
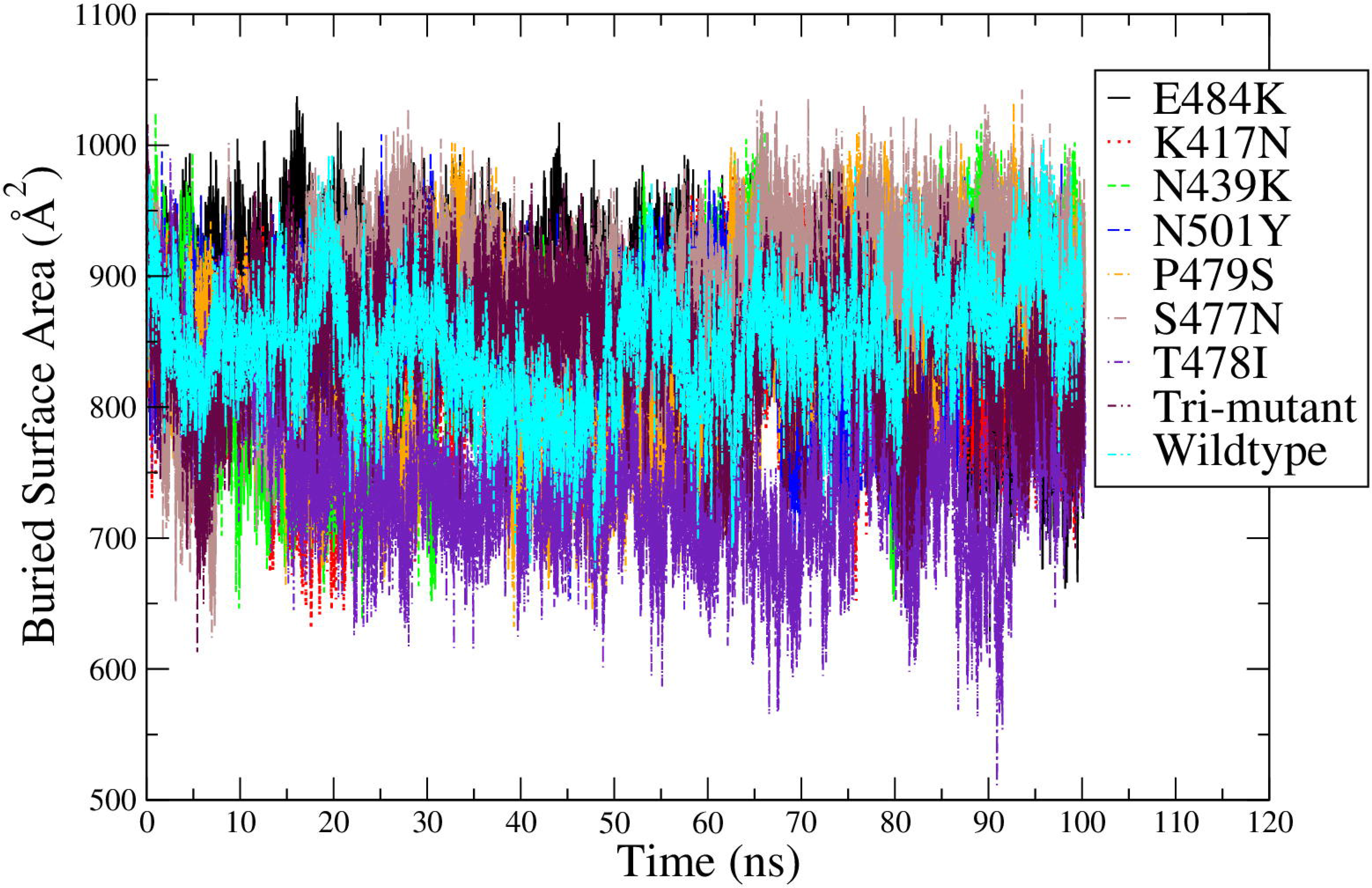
Comparison of the BSA between RBD variants and ACE2 with that of the ACE2&RBD wildtype started from the crystal structure.

#### Distance map result comparison

Calculating the distance map between residues on ACE2 and RBD, the results of RBD-E484K bound with ACE2 in comparison with RBD wildtype bound with ACE2 are shown in Figure 7 as an example. The distance maps of other RBD variants bound with ACE2 are shown in Figure S8 and S9 in SI. Those results show that different RBD variants bound with ACE2 at the interface consistent with that of RBD wildtype and ACE2.

**Figure 7.**
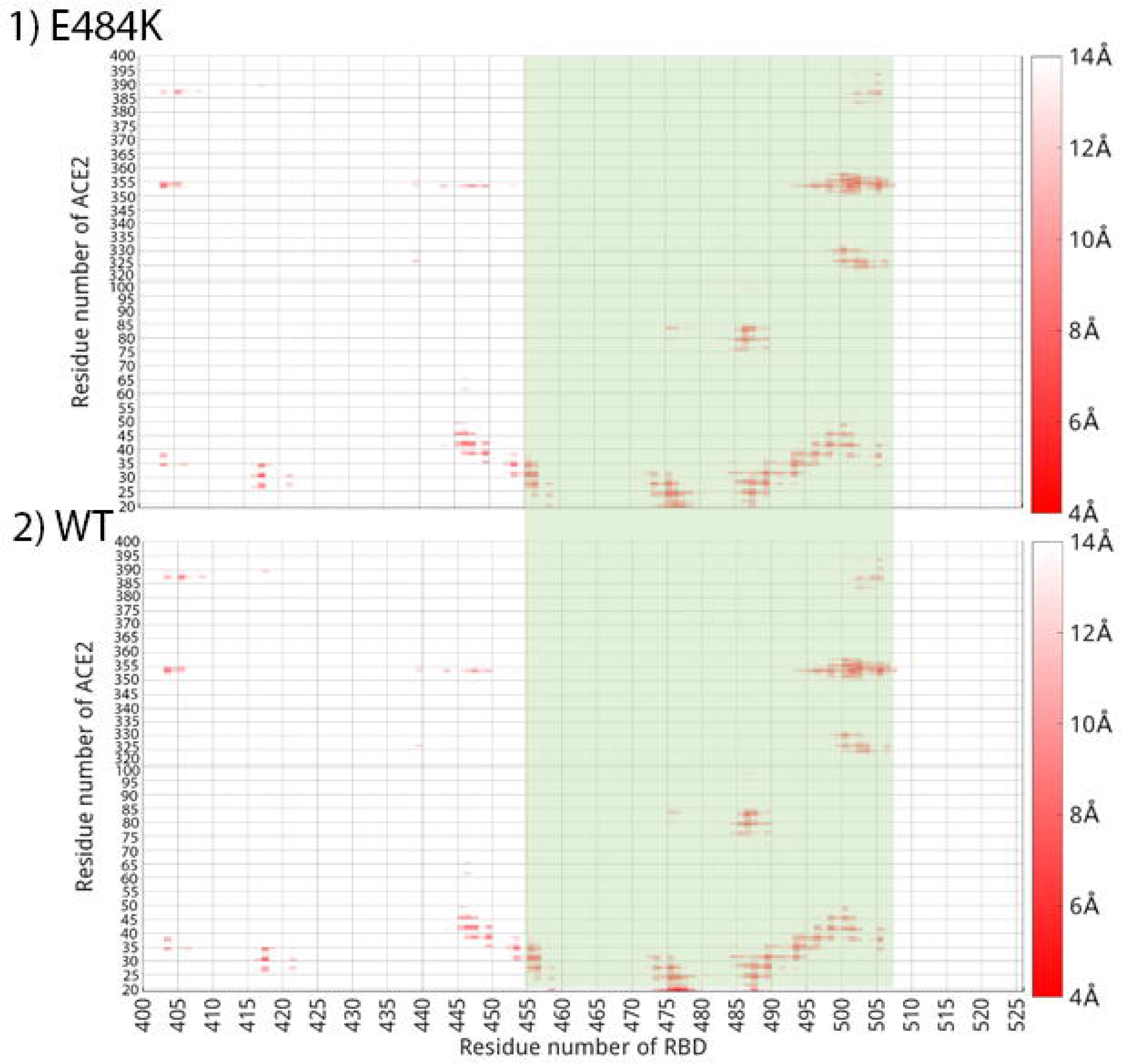
Distance map comparison of 1). RBD-E484K mutant bound with ACE2 and 2). RBD wildtype bound with ACE2. The binding region (RES455 to 505) on RBD are highlighted in light green. The color bars are shown on the right side of the distance maps, with a residue pair distance being no larger than 4 Å shown in red, while a residue pair distance larger or equal to 14 Å shown in white.

Even with the small modification on the residue pairs forming interactions on the binding interface, a similar BSA and distance maps were formed for RBD variants with ACE2 comparing to RBD wildtype. Thus it is not surprising to see that overall the binding structures of RBD mutants&ACE2 complexes are very consistent with the RBD wildtype complex which is the crystal structure of RBD &ACE2 from Lan et al.[51], as the comparison of the initial and final structures for RBD- trimutant&ACE2 shown in Figure 1 (b). The structures of RBD mutants bound with ACE2 showed limited deviation from the original structure, which can agree with RMSD, RMSF, BSA and hydrogen bonds results.

In summary, RBD mutants bind with ACE2 consistently and at a very similar interface. Because the spot mutation did not change the binding interface of RBD and ACE2 significantly, it is reasonable to start from the crystal structure of ACE2&RBD to do FEP simulation and predict the spot mutation effect on ΔΔG.

### 4). FEP result

Analyzing the FEP simulation results, ΔΔG was predicted for RBD mutants. The results in comparison with Upadhyay [39] are shown in Figure 8. The comparison of simulation prediction with three more available literatures [34,35] are shown in Table 2.

**Figure 8.**
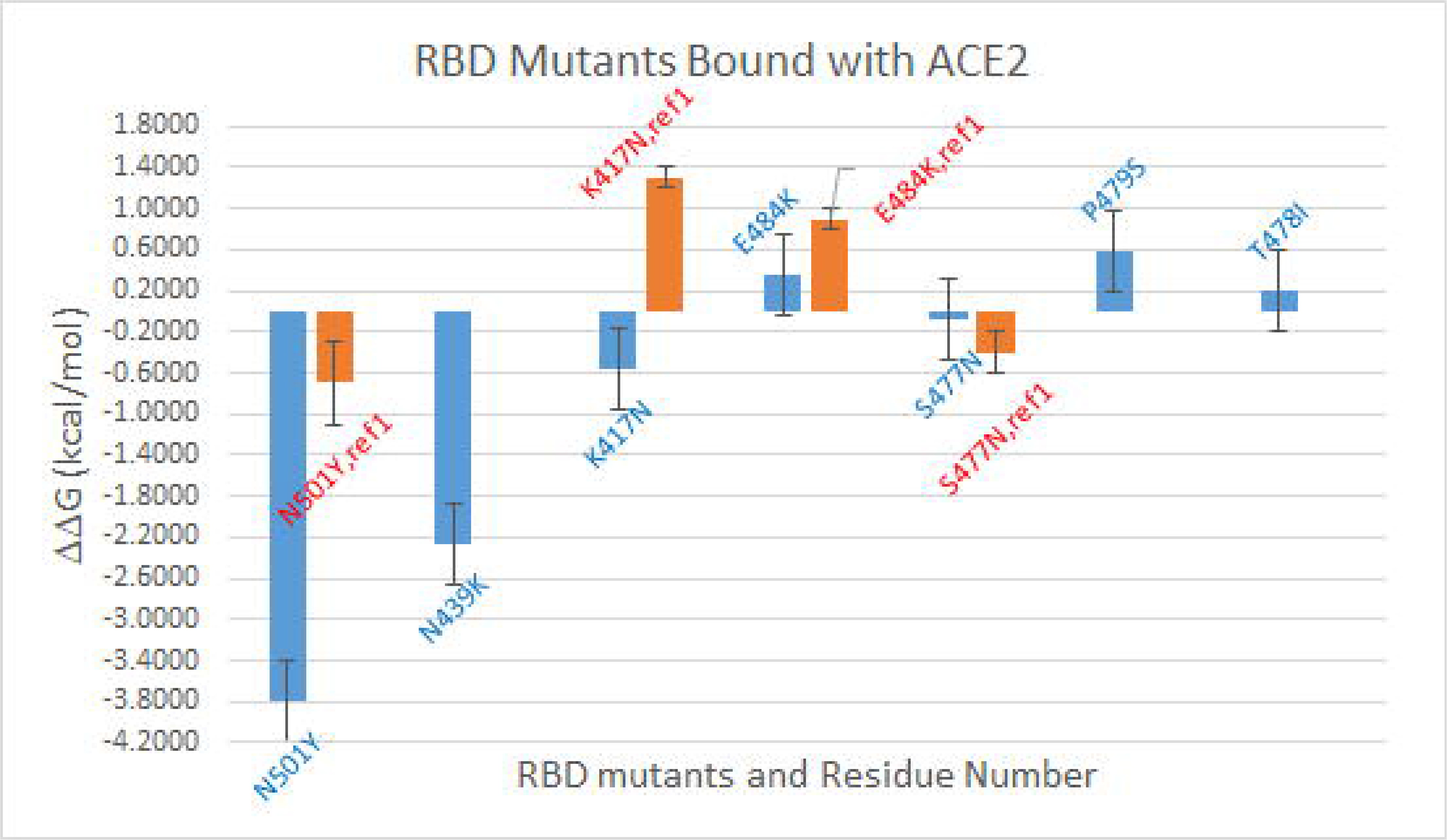
ΔΔG for RBD variants binding with ACE2 predicted from FEP (shown in blue bars) in comparison with literature data from Upadhyay et al. (shown in orange bars). Reference ΔΔG was calculated based on available data in the table as ΔΔG=ΔG_mutant_-ΔG_wt._

**Table 2.** ΔΔG predicted from FEP in comparison with available literature data.

| Mutants | FEP prediction<br>(kcal/mol) | Upadhyay et<br>al. (kcal/mol) | Luan et al.<br>(kcal/mol) | Laffeber<br>(kcal/mol) | Barton et al.<br>(kcal/mol) |
| --- | --- | --- | --- | --- | --- |
| N501Y | -3.80±0.17 | -0.70±0.40 | -0.81 | 1.3 | 1.43±0.04 |
| N439K | -2.26±0.20 |  |  |  |  |
| K417N | -0.56±0.21 | 1.30±0.10 |  | -0.8 | -0.96±0.06 |
| E484K | 0.36±0.27 | 0.90±0.10 |  | 0.2 | 0.20±0.04 |
| S477N | -0.08±0.09 | -0.40±0.20 |  |  | 0.33±0.04 |
| P479S | 0.58±0.07 |  |  |  |  |
| T478I | 0.19±0.08 | 0.3±0.20 |  |  |  |

Based on Figure 8, N501Y, N439K, K417N contribute to the binding of RBD with ACE2, T478I and S477N have a negligible effect, while neither E484K nor P479S contributes to the binding. Upadhyay et al. [39] did ITC experiments on the binding of RBD mutants with ACE2. Calculating ΔΔG using equation: ΔΔG=ΔG_mutant_-ΔG_wt_ based on available data, ΔΔG data from Upadhyay are shown in Figure 8 and Table 2 above. As can be seen from Upadhyay et al., N501Y can increase the binding of RBD with ACE2, T478I has a negligible effect while E484K can make the binding weaker. Our prediction of those three mutations can agree with Upadhyay et al, and the effect of N501Y also agrees with Luan et al. [41] who conducted all-atom molecular dynamics simulations and FEP on RBD-N501Y mutant binding with ACE2, and predicted the ΔΔG=-0.81kcal/mol as shown in Table 2.

However, our FEP predicted that S477N should have a negligible effect on RBD binding with ACE2, while a stronger binding of RBD-S477N variant with ACE2 by Upadhyay et al. Although our prediction deviated from Upadhyay et al, the effect of S477N from Upadhyay et al. was opposite to the binding of RBD and ACE2 by Barton et al. Our FEP prediction on K417N is opposite to Upadhyay et al’s work, although the FEP prediction can agree with the experimental work by Laffeber et al [72] and Barton et al. [73] as shown in Table 2. The DDG result from Laffeber et al. [72] as shown were calculated after converting experimental data K_D_ for wildtype and mutants to DG using the relationship as: DG_exp_=-RTlnK_D_ at T=298K.

N439K, which was found in multiple regions in the world and was a quite common mutation, can enhance the binding of RBD with human ACE2 receptor while evade antibody mediated immunity as found by Thomson et al. [37]. FEP predicted that N439K can decrease the binding free energy of RBD with ACE2 thus should increase the binding affinity between them. Our prediction can agree with Thomson et al.’s result [37].

Based on our FEP prediction, both N501Y and K417N can increase the binding affinity of RBD and ACE2, but E484K has a negative effect on the binding. Because of that, the triple mutants of K417A-E484A-N501A should contribute to the binding of RBD with ACE2 if assuming a linear relationship between the effect of each mutation and the total effect. Because of that, it is expected that ACE2 should bind with the triple mutant stronger than the wildtype. That agrees with the infectivity of Beta variant. The FEP prediction on N501Y suggests that Alpha variant should be more infectious than the original SARS-CoV-2. That also agrees with experimental and clinic findings.

### 4). Dihedral angle correlation covariance result

In order to find out how spot mutations affect the dynamic correlation network of RBD, dihedral angle ϕ, φ combined covariance matrix was calculated for RBD in both wildtype and variant forms, and in both bound and free states. The dihedral angle covariance matrices revealed a set of highly correlated residues in RBD-N501Y and RBD-trimutant, as presented in Figure 9 (Left) and (Right) individually. The RBD wildtype bound with ACE2 covariance matrix is shown in the lower-right triangle below the diagonal line while the covariance matrices of RBD-N501Y variant and tri-mutant variant are shown in the upper-left triangle above the diagonal line in both Figure 9 (Left) and (Right). RBD-N501Y variant lost the original correlation in RBD wildtype between RES501 and other residues as pointed out by the orange arrows, but other correlations in the range of RES470 to RES483 became stronger in RBD-N501Y variant (pointed out by red arrows) which were weak in the original RBD wildtype. Another region around RES370 (pointed out by red arrows) which is also faraway from RES501 site, shows a strong correlation in RBD-N501Y variant, although such kind of strong correlation was not present in the RBD wildtype. Overall, a stronger correlation shows up in RBD-N501Y variant comparing to the RBD wildtype. Similarly, RBD-trimutant (three mutations as N501Y, E484K, K417N) (Figure 9(Right)) has more discrete correlations (pointed out by red arrows) which were not originally in the RBD wildtype, while the original correlations between N501, E484 and K417 and other residues (pointed out by orange arrows) were lost. RES501 region in RBD-tri-mutant is strongly correlated with other regions as pointed out by red arrows.

**Figure 9.**
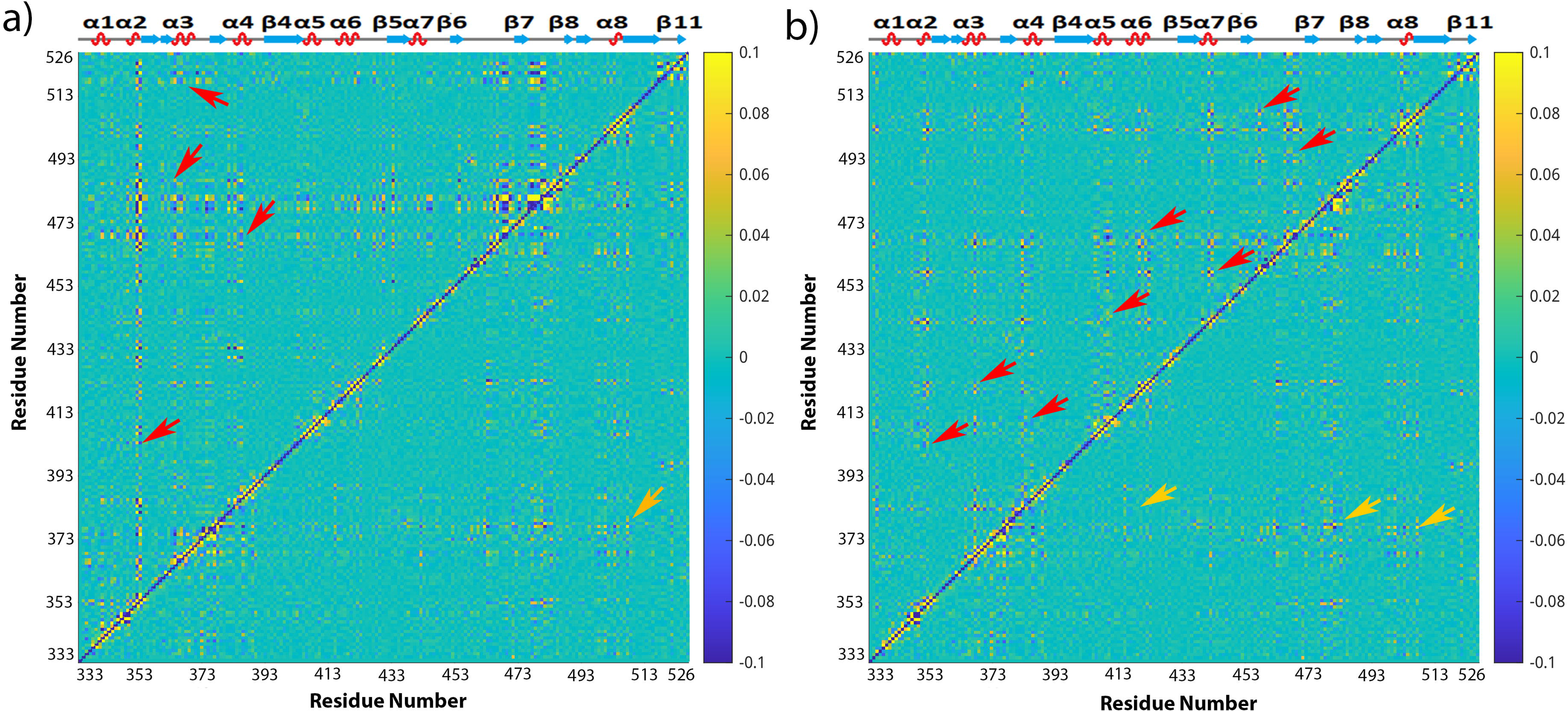
Comparison of the averaged mixed dihedral angle covariance matrix for RBD-N501Y (upper-left triangle above the diagonal line) with RBD wildtype in bound state (lower-right triangle) (Left), and the averaged mixed dihedral angle covariance matrix for RBD-trimutation (upper-left triangle above the diagonal line) in comparison with RBD in wildtype (Right). The color bars of covariance map are shown on the right of the figures from blue to yellow corresponding to the covariance coefficients in the range of -0.1 to 0.1. The secondary structure of RBD is shown on the top of the covariance matrices, and the residue numbers of RBD are labelled on both the x and y-axes.

The correlation matrices for RBD-E484K variant and RBD wildtype in free form were analyzed with results in comparison with RBD wildtype in bound state shown in Figure 10 (Left) and (Right), and results for other RBD variants in Figure S10 to S12. It was interesting to see that RBD-E484K has strong correlation in the head (RES334-374) and tail (RES520-526) regions which are faraway from each other, comparing to the RBD wildtype. The RBD wildtype in free state has a stronger correlation at its tail region but weaker correlation at its head region than the RBD wildtype in bound state. RES372 region has strong correlation with other residues. Since the RBD in free state simulation starts from the crystal structure of RBD (by removing the bound ACE2 in PDB ID of 6M0J) which is in open state, while binding of RBD with ligand can make the binding region more rigid [47], that suggests that binding with ACE2 weakened the correlation of RBD in its tail region which is the binding interface with ACE2.

**Figure 10.**
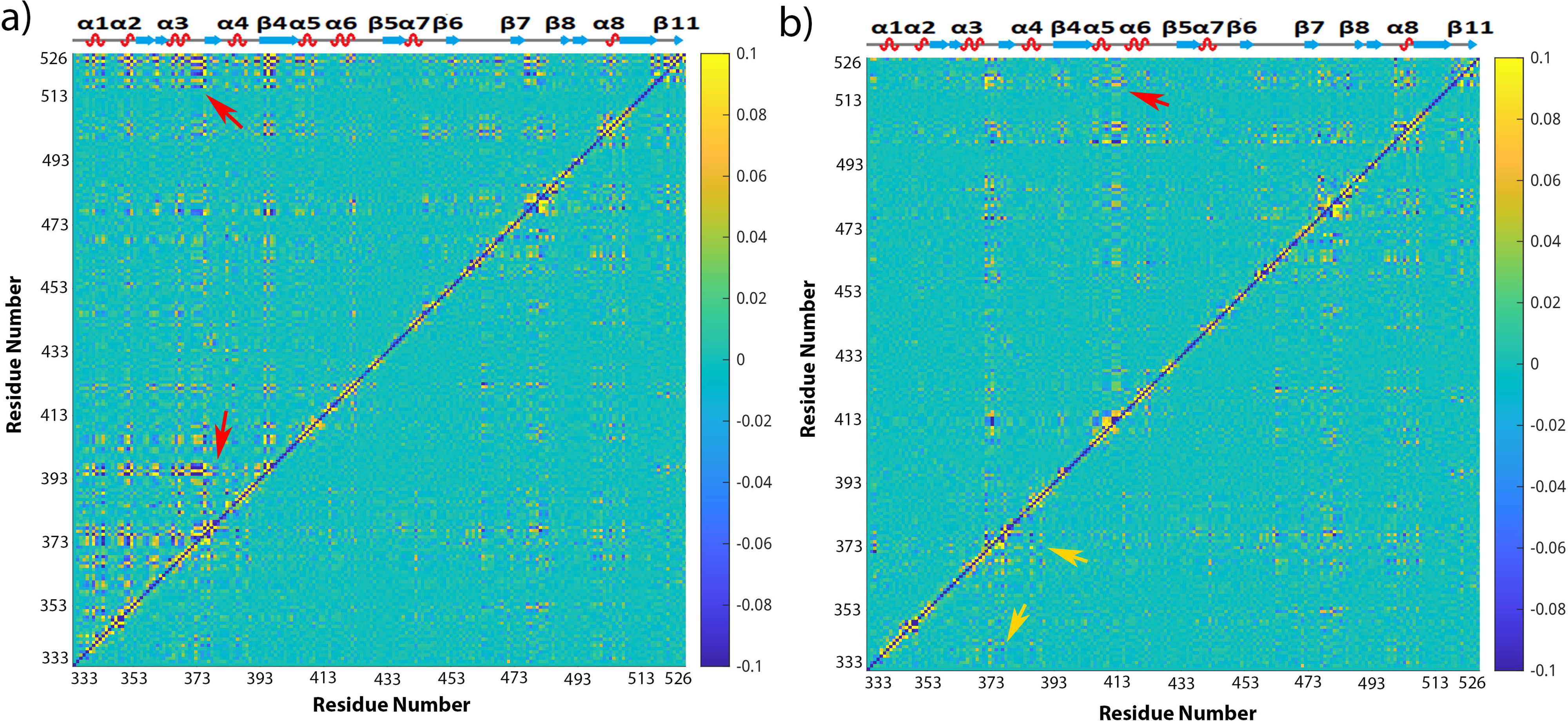
Comparison of the averaged mixed dihedral angle covariance matrices for RBD-E484K (Left) and RBD wildtype in free state (Right) (upper left triangle above the diagonal line) with RBD wildtype in bound state (lower right). The color bars of covariance map are shown on the right of the figures from blue to yellow corresponding to the covariance coefficients in the range of -0.1 to 0.1. The secondary structure of RBD is shown on the top of the covariance matrices, and the residue numbers of RBD are labelled on both the x and y-axes.

Similarly, other RBD variants have their correlation matrix changed comparing to the RBD wildtype as results shown in Figure S10 to S12.

Since RES373 on RBD wildtype in bound form, RES372 on RBD wildtype in free form, RES480 on RBD-N501Y, and RES501 on RBD-trimutant have strong correlation with other residues as shown in Figure 9 and Figure 10, those residues were picked as base for RBD in different simulations correspondingly. Mapping the covariance coefficients of the base residue with other residues on the b-factor of the secondary structure of RBD, the results for RBD wildtype in bound form, RBD wildtype in free form, RBD-N501Y, RBD-trimutant are shown in Figure 11(a), (b), (c), and (d) correspondingly (The secondary structures mapped with covariance coefficients for other RBD variants not shown). In Figure 11, residues having a covariance coefficient higher than 0.05 were labelled, and in red for a positive correlation coefficient while in blue for a negative correlation coefficient. Overall, RBD wildtype in free form, RBD-N501Y, RBD-trimutant all have more correlations than the RBD wildtype in bound form. In RBD-N501Y shown in Figure 11(c), RES480 has strong correlation with P479, G476, F456, S469, N440, I441, T500, Y501, P507, G485, H519, C525, G526, L335, N334. Besides that, other residues labelled in red and blue also have correlation with E484. Checking all the secondary structures mapped with covariance coefficients (picking different base residues) for RBD in different forms, some common correlations exist among residues, such as RES366/363, RES373/372, RES402/403, RES440/441, RES443/444, RES465/464, RES475/476, RES485/484, RES516/517, RES519, RES521/522, RES526/525 in different RBD covariance matrices. Those residues are at the head, binding interface and the tail region, which are mostly α-helices or loops on RBD. Thus it suggests that the helices and loops at the head, binding interface and tail regions have stronger dynamic correlations than the β sheet region.

**Figure 11.**
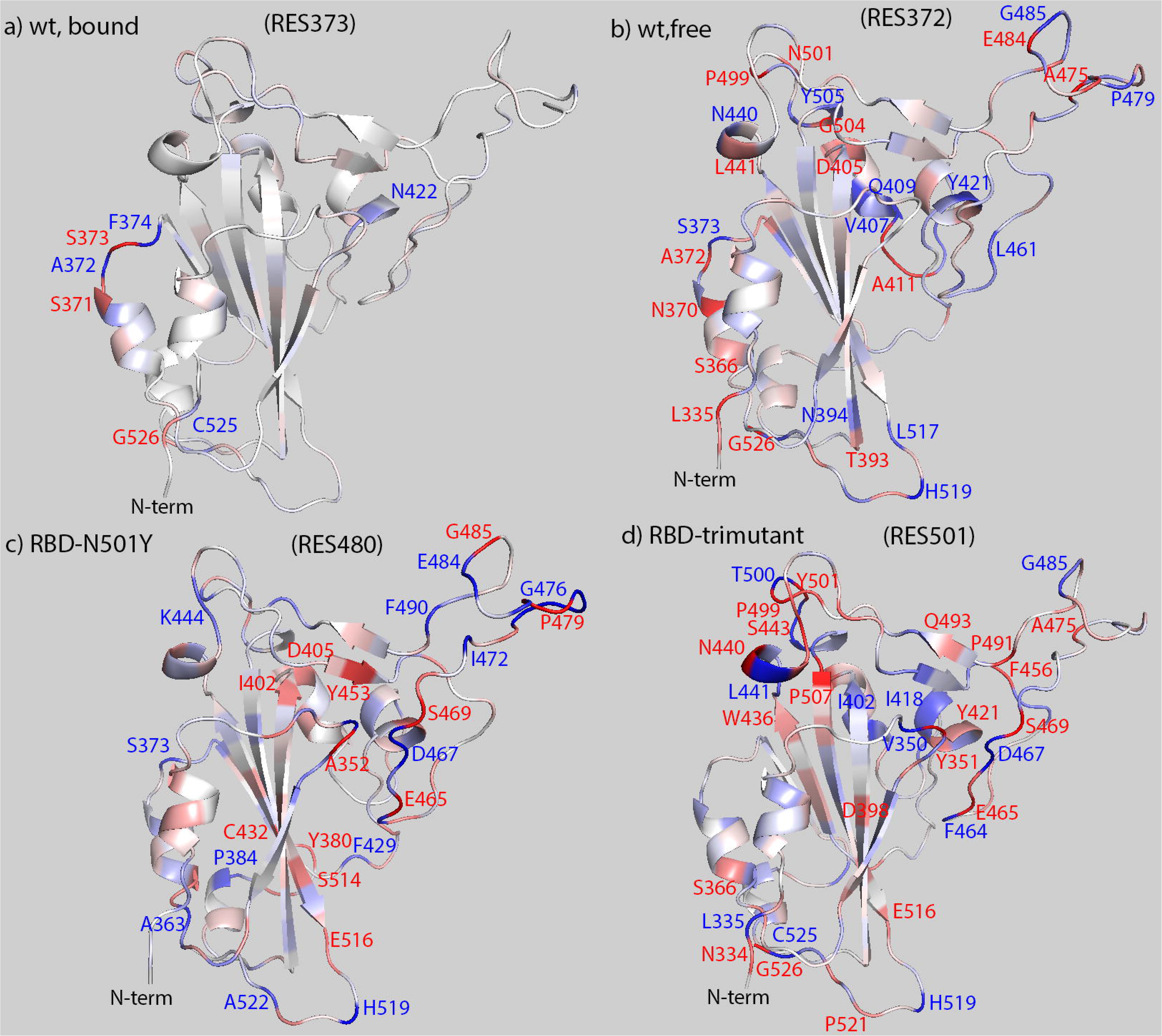
Covariance coefficient between residue 373 (a)/residue 372 (b)/residue 480 (c)/ residue 501(d) and other residues mapped on the structure of RBD wildtype in bound state (a), RBD wildtype in free state (b), and RBD-N501Y variant (c), and RBD-trimutant variant (d). Positive covariance is shown in red while negative covariance in blue and the covariance equals to zero in white. The range of covariance is in the range of -0.1 to 0.1. Residues have positive correlation with the picked residue are labelled in red while in blue for negative correlation.

Because there are interactions between atoms inside a protein, the protein can be viewed as a network. Treating the CA atom in each residue as a node, the interactions between nodes inside the network can be analyzed. Multiple nodes with strong interactions can form one community, which is a subnetwork that partitions the original network in the protein. The communication communities can be calculated using VMD program as explained in the Materials and Methods section. The dynamical network and communication communities in RBD wildtype and variants were analyzed based on residue−residue interactions over time. It was found that usually 7 to 9 network communities formed in RBD in different forms and at different states. The results for RBD-N501Y, RBD wildtype in bound form, RBD wildtype in free form are shown in Figure 12, and in Figure S13 to S14 for other RBD variants. Interestingly, the largest community (shown in blue) always extends from the tail region of RBD to the binding interface in the community maps. The largest community includes the following residues in common: RES335, RES348, RES373, RES428, RES437, RES471, RES490, RES493, RES495, RES496, RES516, RES519, RES521-526. Those residues are at the N-term, central, binding region, and tail of RBD.

**Figure 12.**
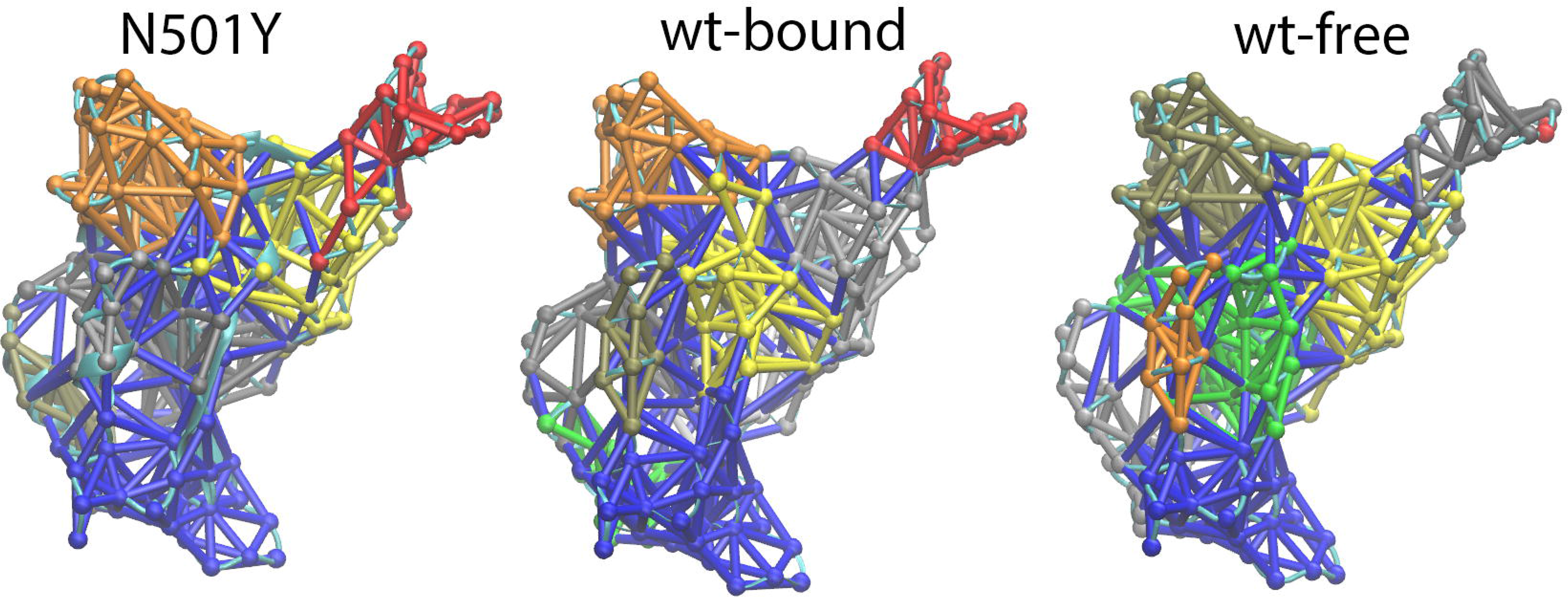
Community networks formed in the RBD-N501Y(Left), in wildtype bound form (middle) and wildtype free form (Right) based on MD simulation and dynamical network analysis. The RBD network structures are oriented in the same direction as the RBD shown in Figure 1.

Such kind of extensive community showed up in all other RBDs as their network community maps shown in Figure 12 and Figure S13 to S14. That suggests that there should be an intrinsic dynamic network community inside RBD distributed from the tail region to the central, and to the binding interface, which is consistent with the dihedral angle covariance result shown in Figure 11. Thus, the dynamics at the RBD binding interface region should be transmitted to the tail of RBD thus to the other sections of S protein. It is expected that disrupting of this intrinsic network can affect the binding of RBD with ACE2 and with antibodies.

## Discussion

Based on the findings from Chan et al. [43], a larger BSA suggests a stronger binding of RBD with ACE2, thus it is expected that E484K, N439K and S477N variants should form a stronger binding with ACE2 than other variants, while the RBD-trimutant should form a less stable binding with ACE2 based on results shown in Figure 6. However, those cannot agree with the ΔΔG result predicted from FEP as shown in Figure 8, which shows that N439K mutation should decrease the binding free energy of RBD and ACE2, but not E484K. S477N mutation almost does not change the binding free energy of RBD and ACE2. Thus, the size of BSA does not correlate to the binding affinity of RBD and ACE2 based on our test in this project.

Up to now, different results on spot mutation effects on the binding affinity of RBD with ACE2 were reported. Tian et al. [74] found that N501Y mutation can increase the binding affinity of RBD with ACE2 receptor using both experimental and simulation methods, thus a higher rate of transmission of SARS-CoV-2 variant. However, they found that E484K did not contribute to the binding of RBD with ACE2. That is different from Wang et al. [75], who found that E484K mutant on RBD can increase the binding affinity of RBD with ACE2 receptor, while decrease the binding affinity of RBD with antibodies. Upadhyay et al. [39] found E484K had a weaker binding affinity to ACE2 than the wildtype, although N501Y had a stronger binding affinity to ACE2 and S477N and T478I have a similar affinity to ACE2. Based on experimental test result, Wang et al. [76] suggested that T478I and N501Y mutants can increase the infectivity of SARS-CoV-2, and can potentially facilitate the transmission of SARS-CoV-2 virus to many other animal hosts. Therefore, more attention should be paid to SARS-CoV-2 variants with these two mutations. In this project, it has been found that spot-mutation on RBD do not change the binding interface between ACE2 and RBD comparing to the wildtype. However, each spot-mutation can make different contributions to the binding free energy of RBD with ACE2. N501Y can increase the binding of RBD with ACE2 significantly based on FEP prediction. That agrees with findings from some other researchers as shown in Table 2. Such kind of discrepancy in experimental result emphasized the importance of theoretical prediction and systematical comparative study on the SARS-CoV-2 spot-mutation variants.

Based on the dihedral angle covariance matrix calculation, it was found that there are some common correlations in RBD, no matter in variant or wildtype form, in bound or free state. Those residues are mostly located in the largest network community predicted by dynamic network analysis with results shown in Figure 12 and Figure S13 to S14. Since the community is intrinsic in RBD, and it extends from tail region of RBD to the binding interface region, it is expected that mutation on those residues can affect the binding of RBD with ACE2 and with antibodies.

Since the spike protein of SARS-CoV-2 is a trimer and each monomer has more than 1000 residues [77], more work should be done in order to understand how the spot mutations on S protein affect its structure, dynamics and function thus to supply insight on designing novel drugs against future pandemic.

## Conclusions

In this project, all-atom molecular dynamics simulations using NAMD program were conducted on seven RBD single mutants and one RBD triple mutant binding with ACE2. It was found that spot mutation on RBD does not affect the binding interface of RBD with ACE2 significantly. Similar number of hydrogen bonds still formed on the binding interface between RBD and ACE2 after the spot mutation, although some of the exact residues forming hydrogen bonds changed. The distance map result shows that RBD mutant and ACE2 form the binding interface consistent with the RBD wildtype, and they also form a similar buried surface area on the interface. Thus, the structures of RBD mutants bound with ACE2 are mostly consistent with RBD wildtype with ACE2. Calculating ΔΔG of spot mutation using FEP method, it was found that N501Y, N439K, K417N can improve the binding of RBD with ACE2; E484K and P479S could not increase the binding of RBD with ACE2; S477N and T478I do not affect the binding of RBD with ACE2. The spot mutation changed the dynamic correlation network of residues on RBD based on the dihedral correlation calculation, but an intrinsic network community exists in RBD which extends from the tail region to the central region, then to the binding interface region. The result in this project can supply the methodology and molecular insight on studying the molecular structure and dynamics of possible future pandemics and design novel drugs.

## Acknowledgements

This work was supported by National Science Foundation (NSF) Grant NO. 2200138. The simulations were performed using supercomputer time from XSEDE via an award to Zhang L. (MCB160041), and with some trajectory analysis partially using the high performance computers at UTC, and one simulation performed on Unity computers of URI.

## Notes

### Competing Interest Statement

The authors have declared no competing interest.

### Summary of Updates

Some small grammar errors were fixed.

