## supplemental-file for "Structure, Dynamics and Free Energy Studies on the Effect of Spot Mutations on SARS-CoV-2 Spike Protein Binding with ACE2 Receptor"

### Tables

Table S1. Major spot mutations on different SARS-CoV-2 variants in the RBD, and two extra mutations on RBM studied in this work. Reference [29] is listed in the main context.

| Mutation Sites | Alpha | Beta | Gamma | Delta | Omicron |
|---|---|---|---|---|---|
| K417N | | √ | | | √ |
| N440K | | | | | √ |
| G446S | | | | | √ |
| L452R | | | | √ | |
| S477N | | | | | √ |
| T478K | | | | √ | √ |
| E484A | | | | | √ |
| E484K | | √ | √ | | |
| Q493K | | | | | √ |
| G496S | | | | | √ |
| Q498R | | | | | √ |
| N501Y | √ | √ | √ | | √ |
| Y505H | | | | | √ |
| N439K | Happened with high frequency based on genome data from Chen et al. [29] | | | | |
| P479S | Happened with high frequency based on genome data from Chen et al. [29] | | | | |

Table S2. The list of simulations conducted in this project, and the details of the simulations.

| ACE2 bound with RBD mutant | Starting structure | Box size (Å$^3$) | Simulation length (ns) |
|---|---|---|---|
| #1,E484K | Lan et al.[23] | 85.1x95.2x126.5 | 100 |
| #2,N439K | Lan et al.[23] | 85.1x95.2x126.6 | 100 |
| #3,N501Y | Lan et al.[23] | 85.1x95.2x126.6 | 100 |
| #4,P479S | Lan et al.[23] | 85.1x95.2x126.6 | 100 |
| #5,T478I | Lan et al.[23] | 85.1x95.2x127.2 | 100 |

| | | | |
|---|---|---|---|
| #6,K417N | Lan et al.[23] | 85.1x95.2x126.5 | 100 |
| #7,S477N | Lan et al.[23] | 85.1x95.2x127.1 | 100 |
| #8,N501Y-E484K-K417N | Lan et al.[23] | 85.1x95.2x126.9 | 100 |
| #9, RBD wildtype | Lan et al.[23] | 65.7x73.3x74.3 | 100 |
| #10, ACE2 | Lan et al.[23] | 93.3x94.8x78.8 | 100 |
| FEP simulations, | | | |
| Mutant site | Starting Structure | Box size (Å$^3$) | Simulation length (ns) |
| #1,E484K | Lan et al.[23] | 82.4x92.2x134.6 | 3 |
| | | 62.3x70.6x77.8 | 3 |
| #2,N439K | Lan et al.[23] | 82.4x92.2x134.9 | 3 |
| | | 64.2x70.6x78.1 | 3 |
| #3,N501Y | Lan et al.[23] | 82.5x92.3x134.7 | 3 |
| | | 64.4x70.7x78.1 | 3 |
| #4,K417N | Lan et al.[23] | 82.4x92.2x134.8 | 3 |
| | | 64.3x70.6x77.8 | 3 |
| #5,S477N | Lan et al.[23] | 82.4x92.1x134.9 | 3 |
| | | 64.3x70.6x77.9 | 3 |
| #6,P479S | Lan et al.[23] | 82.5x92.3x134.6 | 3 |
| | | 64.2x70.6x77.8 | 3 |
| #7,T478I | Lan et al.[23] | 82.5x92.3x134.6 | 3 |
| | | 64.3x70.6x77.9 | 3 |

Table S3. The residue number and occupancy of hydrogen bonds formed in RBD mutant binding with ACE2 in 100 ns NAMD simulations. Only hydrogen bonds with an occupancy higher than 10% are shown. The residues pairs which form hydrogen bonds on ACE2 and RBD wildtype are highlighted in blue in the ACE2 and RBD mutant jobs.

| Bound with ACE2 jobs | RES#(RBD)-RES#(ACE2) | Occupancy (%) | Bound with ACE2 jobs | RBD-ACE2 | Occupancy (%) |
|---|---|---|---|---|---|
| RBD-K417N | THR500-ASP355 | 42.36 | RBD-N439K | LYS417-ASP30 | 45.3 |
| | GLY502-LYS353 | 38.55 | | GLY502-LYS353 | 43.4 |
| | ASN487-TYR83 | 36.28 | | ASN487-TYR83 | 38.2 |
| | TYR505-GLU37 | 24.61 | | THR500-ASP355 | 32.8 |
| | GLN493-GLU35 | 23.32 | | TYR505-GLU37 | 30.9 |
| | TYR449-ASP38 | 15.63 | | GLN493-GLU35 | 27.6 |
| Bound with ACE2 jobs | RBD-ACE2 | Occupancy (%) | Bound with ACE2 jobs | RBD-ACE2 | Occupancy (%) |
| RBD-T478I | LYS417-ASP30 | 42.9 | RBD-N501Y | GLY502-LYS353 | 54.3 |
| | GLY502-LYS353 | 40.3 | | LYS417-ASP30 | 44.4 |
| | ASN487-TYR83 | 35.8 | | ASN487-TYR83 | 41.7 |
| | GLN493-GLU35 | 10.2 | | TYR505-GLU37 | 31.7 |
| | | | | THR500-ASP355 | 30.0 |
| | | | | GLN493-GLU35 | 22.0 |

| | | | | | |
|---|---|---|---|---|---|
| | | | | THR500-TYR41 | 13.7 |
| | | | | GLN498-GLN42 | 10.2 |
| Bound with ACE2 jobs | RBD-ACE2 | Occupancy (%) | Bound with ACE2 jobs | RBD-ACE2 | Occupancy (%) |
| RBD-P479S | LYS417-ASP30 | 52.4 | RBD-S477N | LYS417-ASP30 | 46.2 |
| | GLY502-LYS353 | 46.2 | | GLY502-LYS353 | 44.9 |
| | TYR505-GLU37 | 39.2 | | ASN487-TYR83 | 40.0 |
| | ASN487-TYR83 | 37.8 | | TYR505-GLU37 | 33.1 |
| | GLN493-GLU35 | 26.1 | | GLN493-GLU35 | 25.7 |
| | THR500-TYR41 | 17.3 | | THR500-ASP355 | 23.6 |
| | TYR449-ASP38 | 14.6 | | THR500-TYR41 | 17.0 |
| | GLN498-LYS353 | 10.9 | | GLN493-LYS31 | 12.5 |
| | | | | GLN498-GLN42 | 11.9 |
| Bound with ACE2 jobs | RES#(RBD)-RES# (ACE2) | Occupancy (%) | Bound with ACE2 jobs | RBD-ACE2 | Occupancy (%) |
| RBD-E484K | LYS417-ASP30 | 40.6 | RBD -N501Y-E484K-K417N | GLY502-LYS353 | 52.0 |
| | ASN487-TYR83 | 40.3 | | ASN487-TYR83 | 33.8 |
| | TYR505-GLU37 | 31.9 | | THR500-ASP355 | 25.6 |
| | GLY502-LYS353 | 30.8 | | GLN493-GLU35 | 23.2 |
| | THR500-ASP355 | 30.7 | | TYR505-GLU37 | 19.1 |
| | GLN498-GLN42 | 21.5 | | THR500-TYR41 | 11.8 |
| | GLN493-GLU35 | 17.6 | | | |
| | GLN498-LYS353 | 17.4 | | | |
| | TYR449-ASP38 | 17.4 | | | |
| | THR500-TYR41 | 13.4 | | | |
| | GLN498-GLN42 | 12.3 | | | |
| Bound with ACE2 jobs | RES#(RBD)-RES# (ACE2) | Occupancy (%) | | | |
| RBD-wt | GLY502-LYS353 | 42.05 | | | |
| | ASN487-TYR83 | 41.46 | | | |
| | LYS417-ASP30 | 34.23 | | | |
| | GLN498-LYS353 | 27.21 | | | |
| | TYR449-ASP38 | 24.67 | | | |
| | TYR505-GLU37 | 22.53 | | | |
| | THR500-TYR41 | 22.35 | | | |
| | GLN493-GLU35 | 21.14 | | | |
| | GLN498-GLN42 | 19.43 | | | |

**Figures**

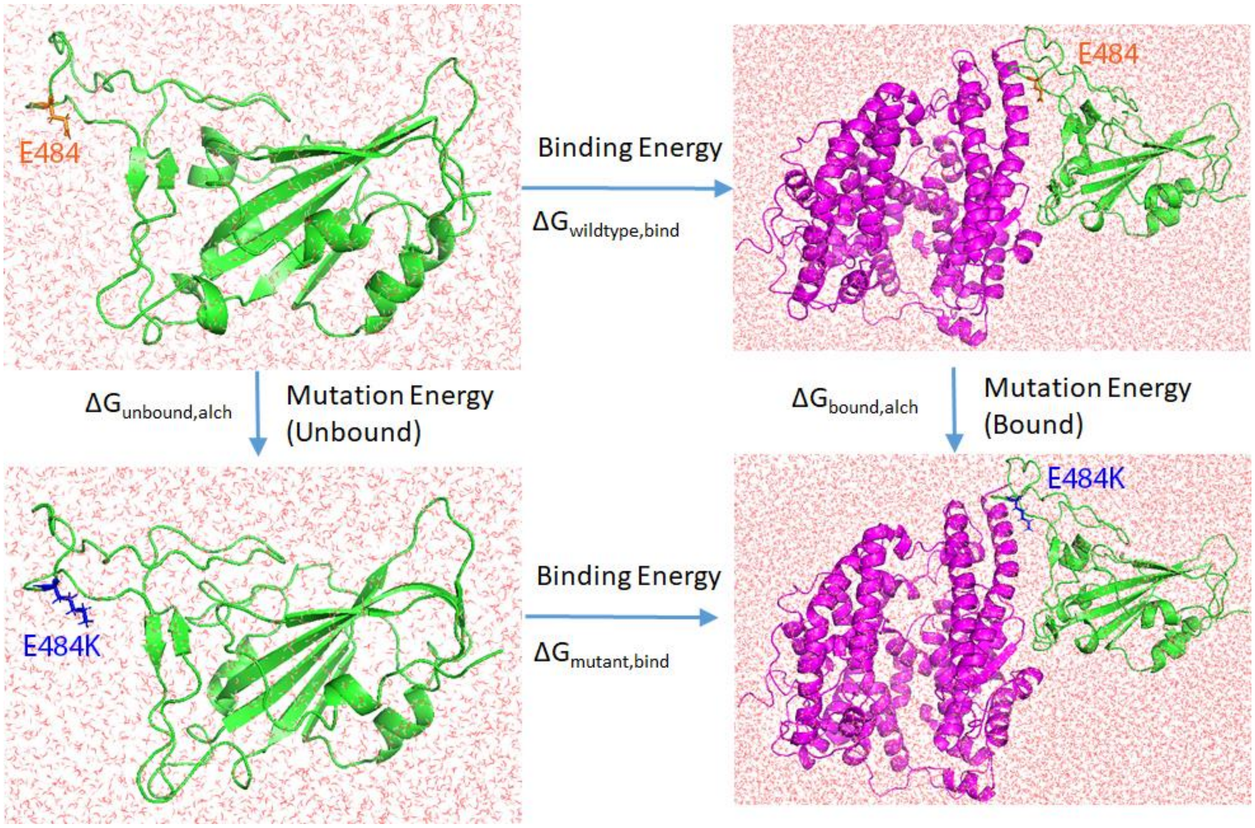


Figure S1. Thermodynamics cycle of FEP calculation, with E484 as the example residue, which is shown in orange sticks. After mutating E484 into K484, it is shown in blue sticks. The RBD is shown in green cartoon while ACE2 shown in magenta cartoon. Water molecules are shown in red sticks and ions are not shown.

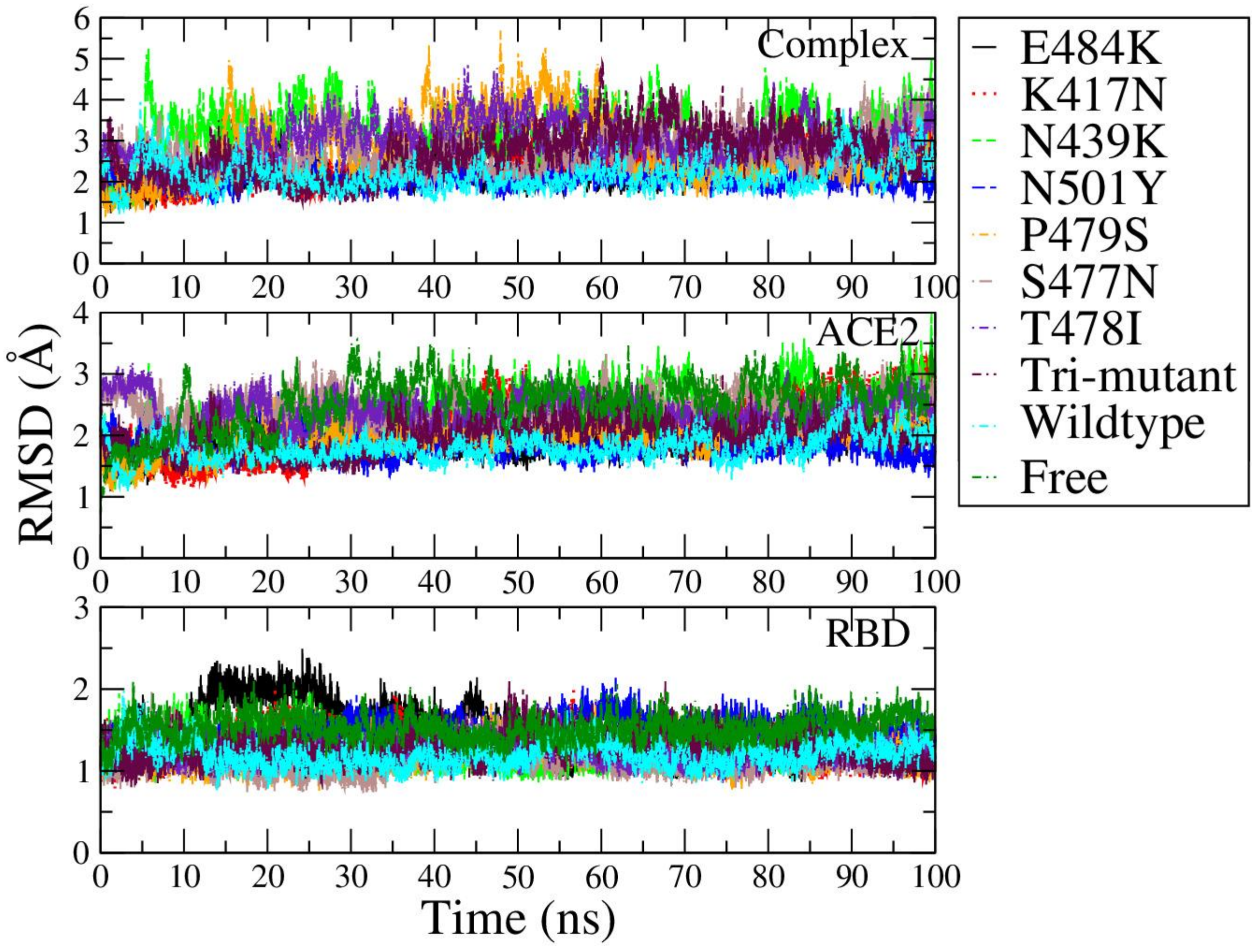


Figure S2. RMSD of ACE2&RBD mutant complexes (Top row), ACE2 (Middle row) and RBD (Bottom row) during 100 ns all-atom NAMD simulations.

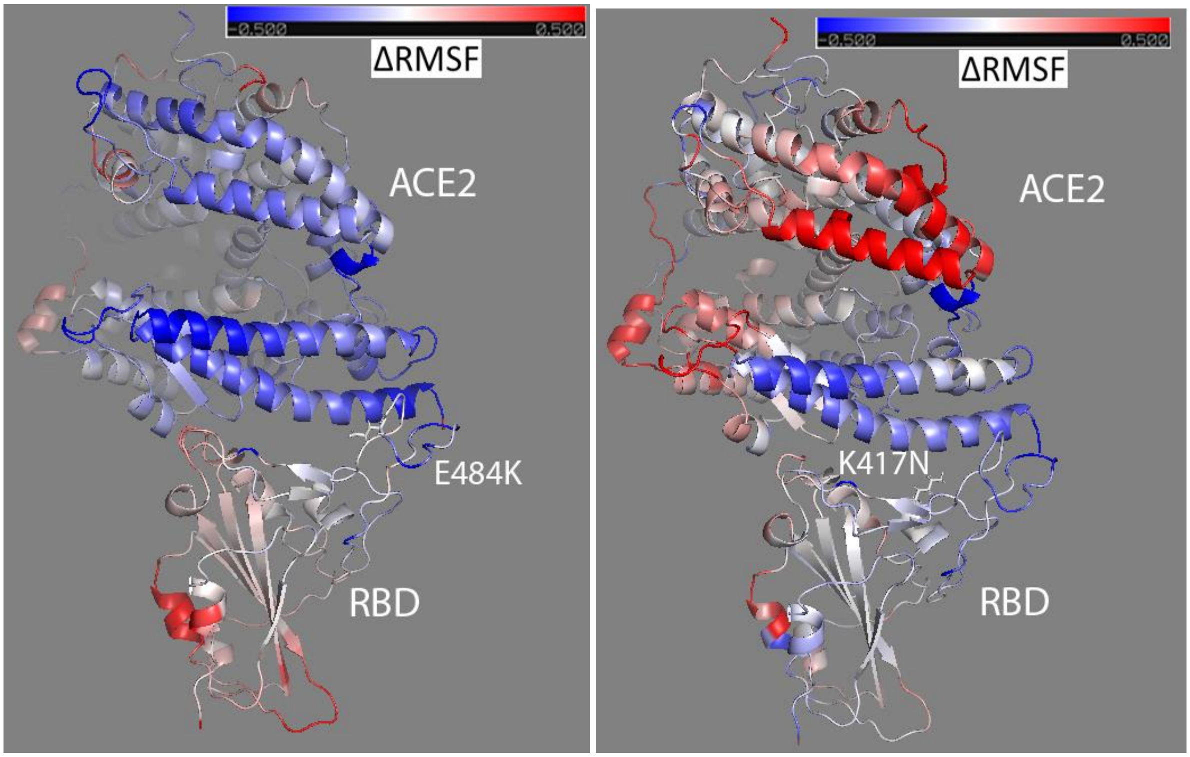


Figure S3. The ΔRMSF of RBD-E484K and ACE2 (Left) from RBD wildtype to variant forms mapped to the initial bound structure of ACE2&RBD-E484K; and the ΔRMSF of RBD-K417N and ACE2 (Right) mapped to its initial bound structure of the complex. Colors range from blue to white to red (with white representing no change in RMSF, blue representing RMSF decrease, and red representing RMSF increase). The ΔRMSF is in the range from -0.5 (totally blue) to 0.5 (totally red).

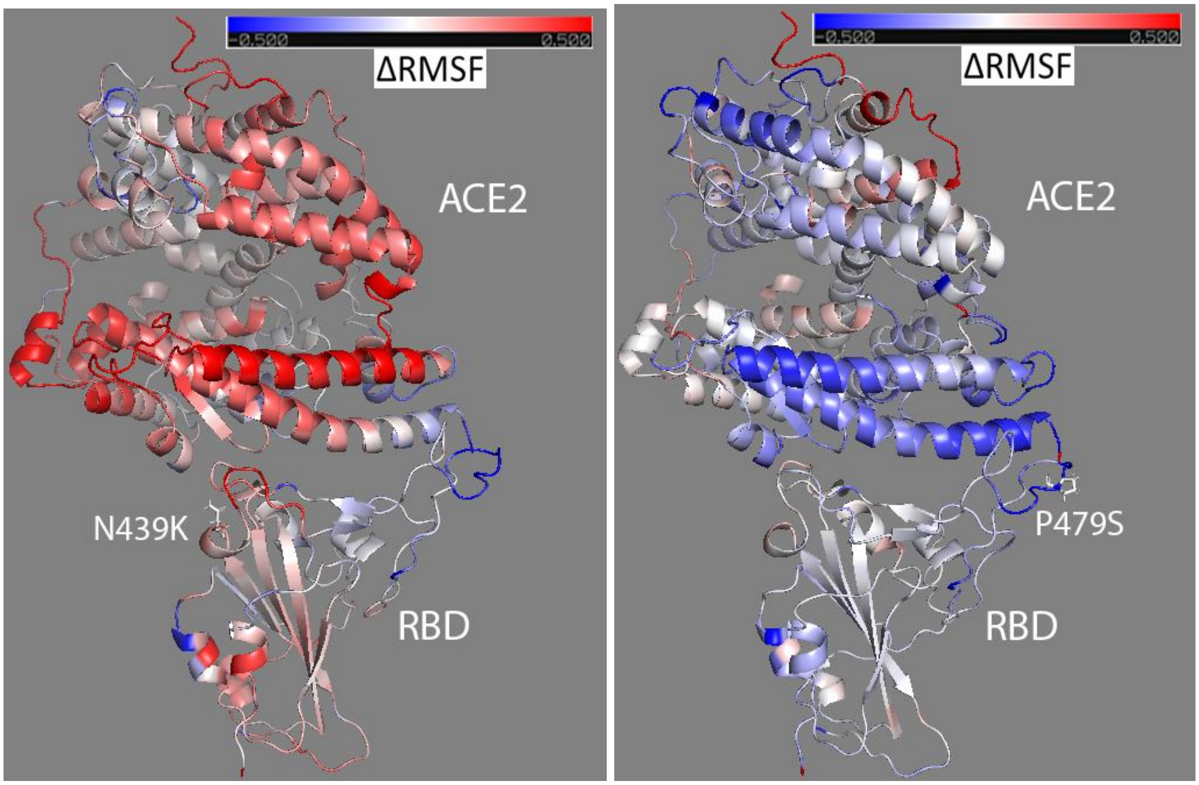


Figure S4. The ΔRMSF of RBD-N439K and ACE2 (Left) from RBD wildtype to variant forms mapped to the initial bound structure of ACE2&RBD-N439K; and the ΔRMSF of RBD-P479S and ACE2 (Right) mapped to its initial bound structure of the complex. Colors range from blue to white to red (with white representing no change in RMSF, blue representing RMSF decrease, and red representing RMSF increase). The ΔRMSF is in the range from -0.5 (totally blue) to 0.5 (totally red).

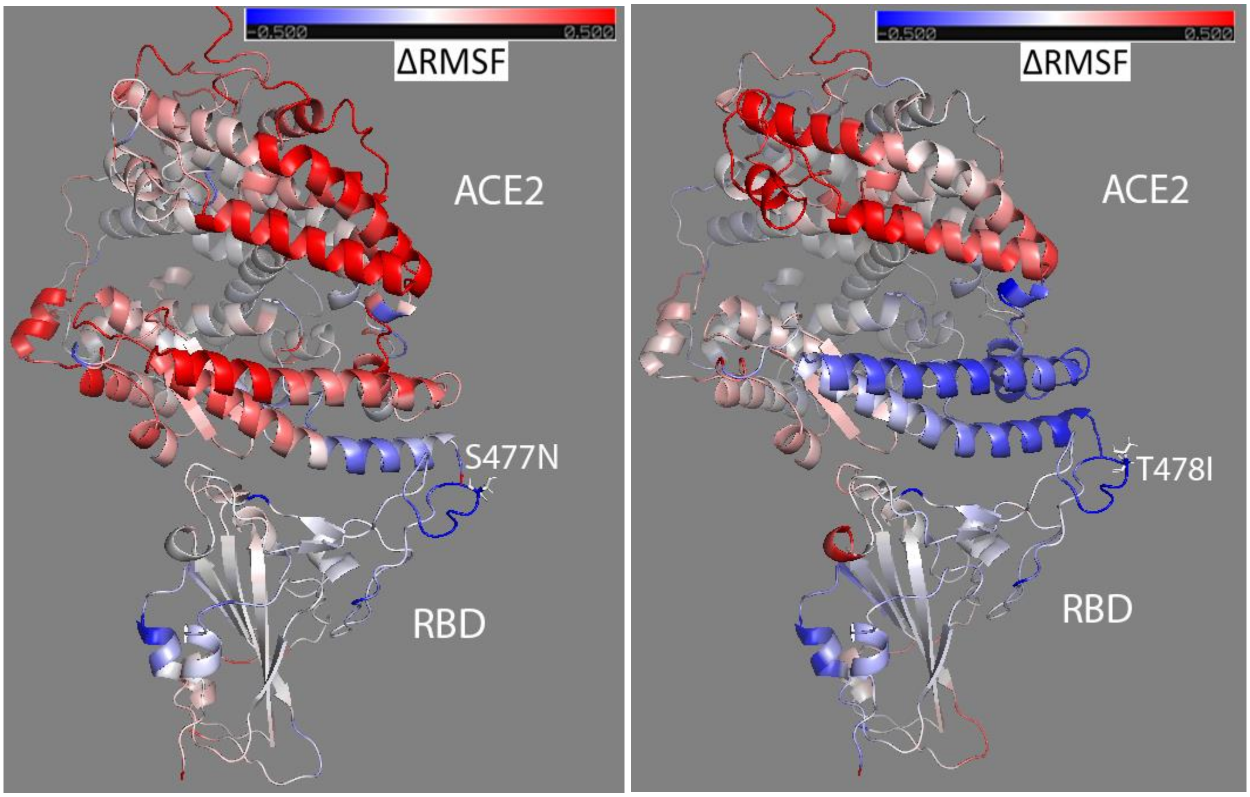


Figure S5. The ΔRMSF of RBD-S477N and ACE2 (Left) from RBD wildtype to variant forms mapped to the initial bound structure of ACE2&RBD-S477N; and the ΔRMSF of RBD-T478I and ACE2 (Right) mapped to its initial bound structure of the complex. Colors range from blue to white to red (with white representing no change in RMSF, blue representing RMSF decrease, and red representing RMSF increase). The ΔRMSF is in the range from -0.5 (totally blue) to 0.5 (totally red).

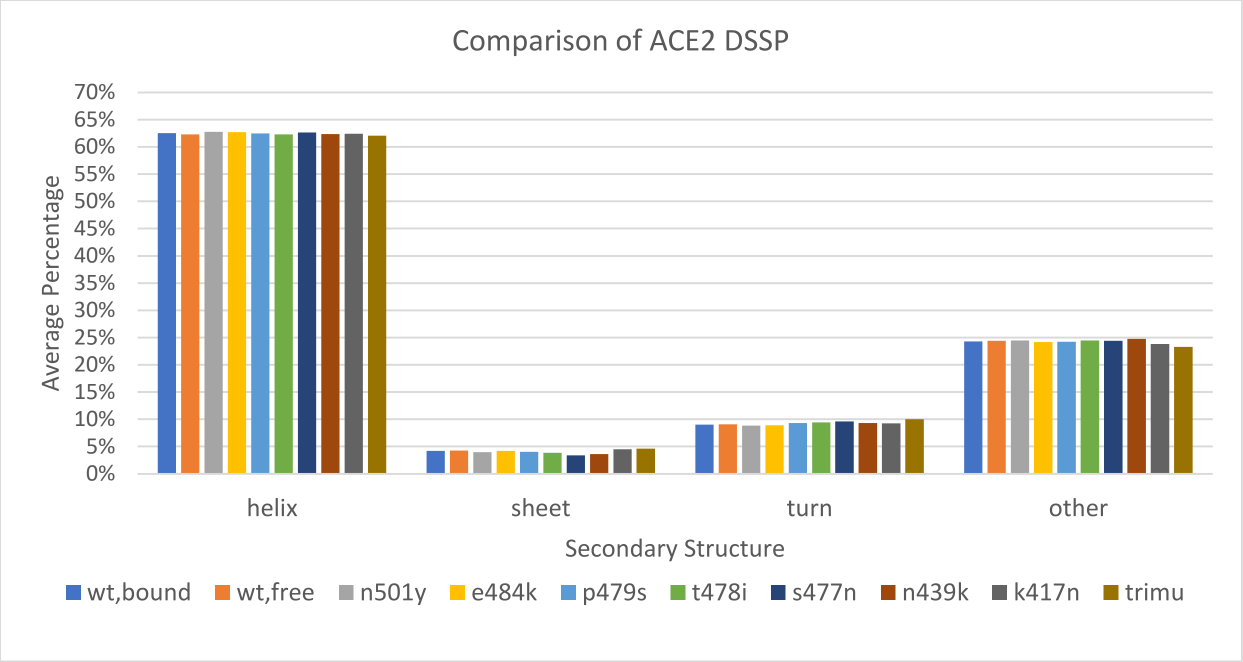


Figure S6. Comparison of ACE2 DSSP from different simulations.

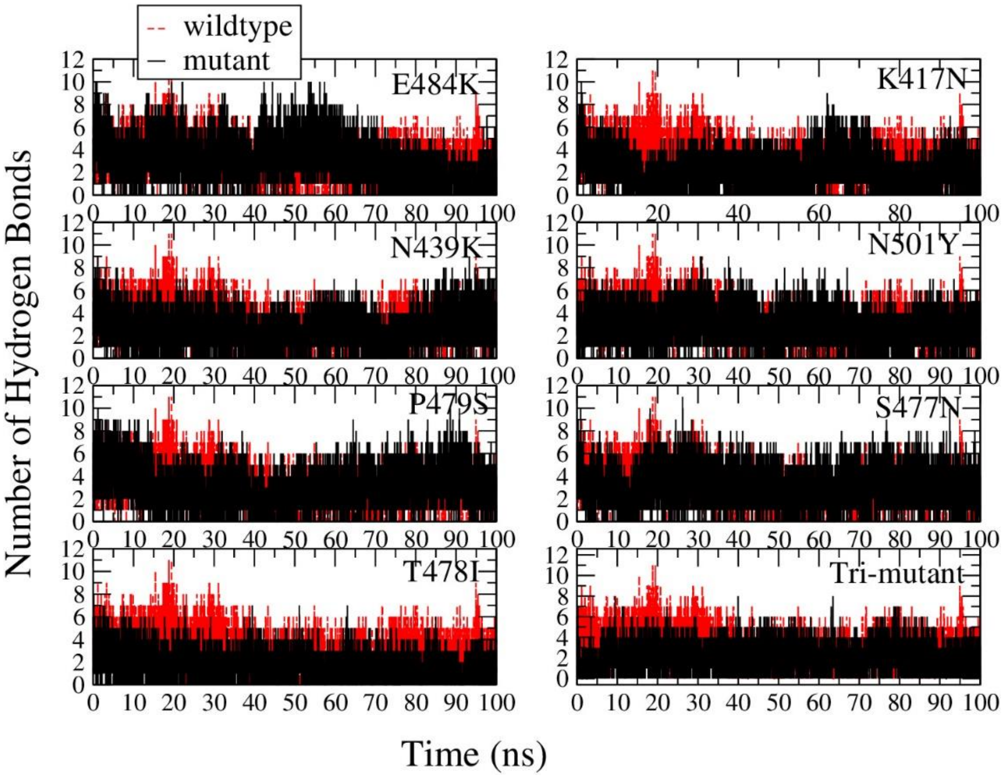


Figure S7. The number of hydrogen bonds comparison between RBD mutants bound with ACE2 and the wildtype from simulations started from wt-bound simulations. The result from mutants are shown in black in front, while red for the wildtype in back.

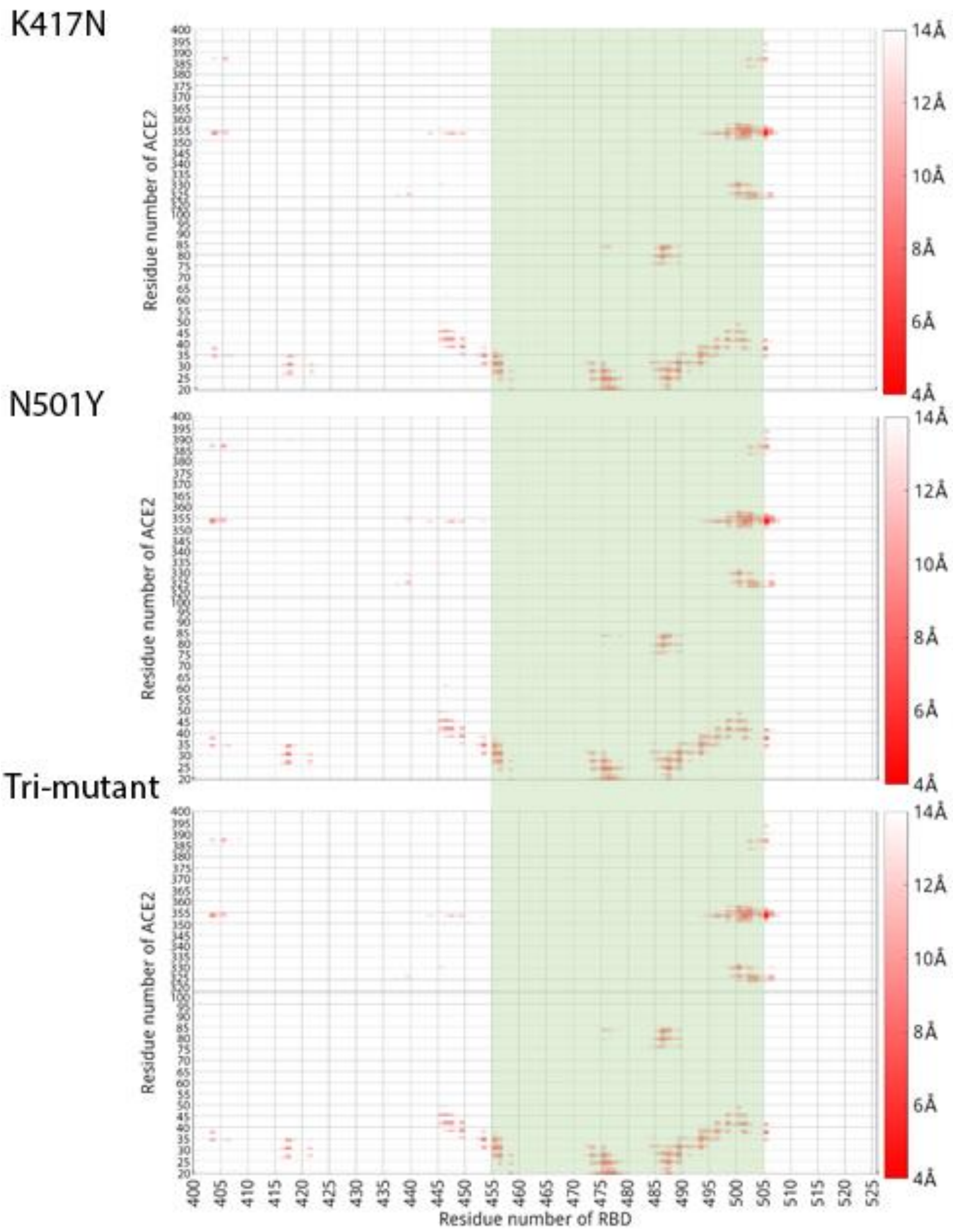


Figure S8. The distance map of RBD-K417N, RBD-N501Y, RBD-trim-mutant variants binding with ACE2. The binding region (RES455 to 505) on RBD are highlighted in light green. The color bars are shown on the right side of the distance maps, with a residue pair distance being no larger than 4 Å shown in red, while a residue pair distance larger or equal to 14 Å shown in white.

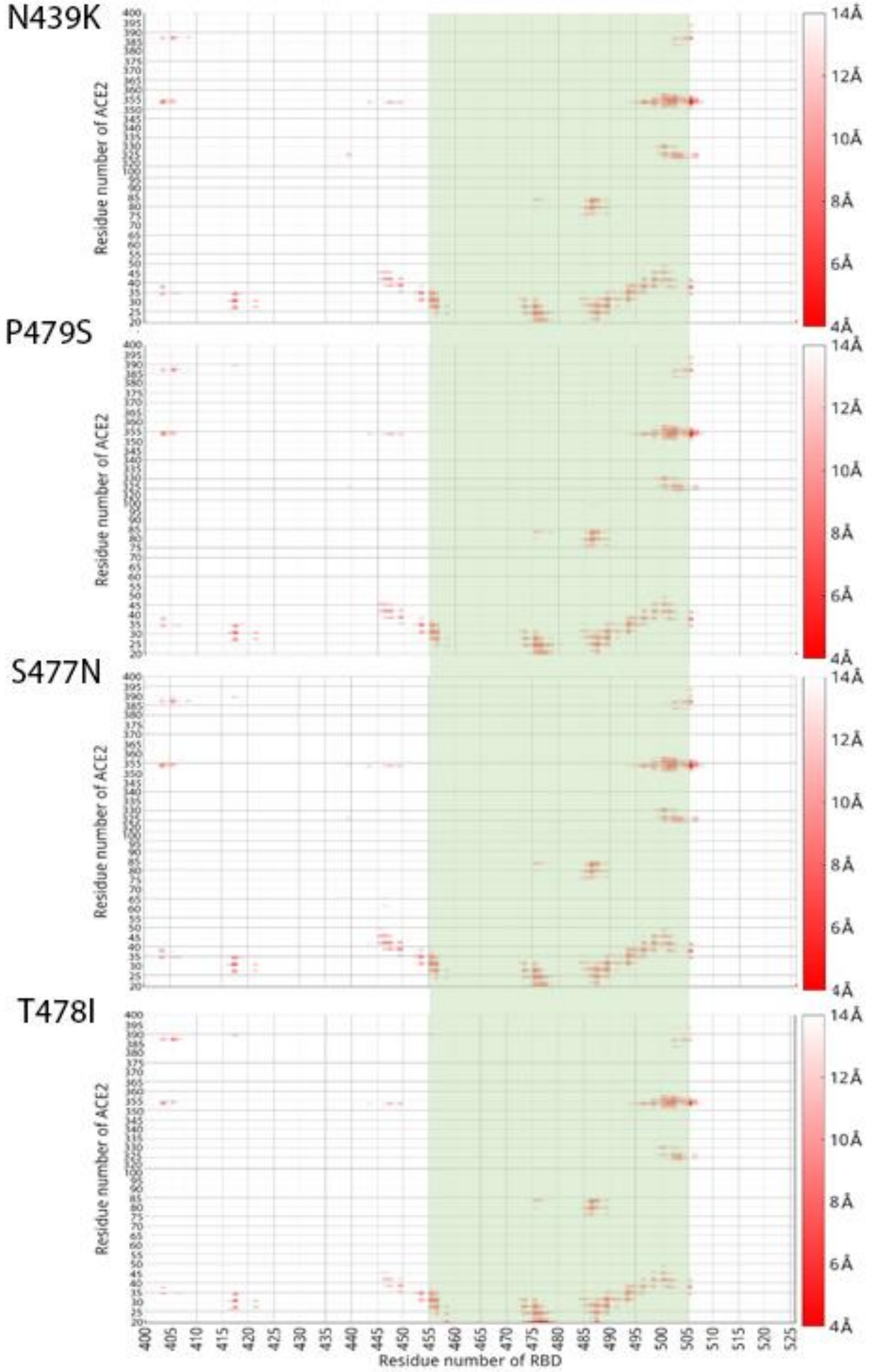


Figure S9. The distance maps of RBD-N439K, RBD-P479S, RBD-S477N, RBD-T478I variants binding with ACE2. The binding region (RES455 to 505) on RBD are highlighted in light green. The color bars are shown on the right side of the distance maps, with a residue pair distance being no larger than 4 Å shown in red, while a residue pair distance larger or equal to 14 Å shown in white.

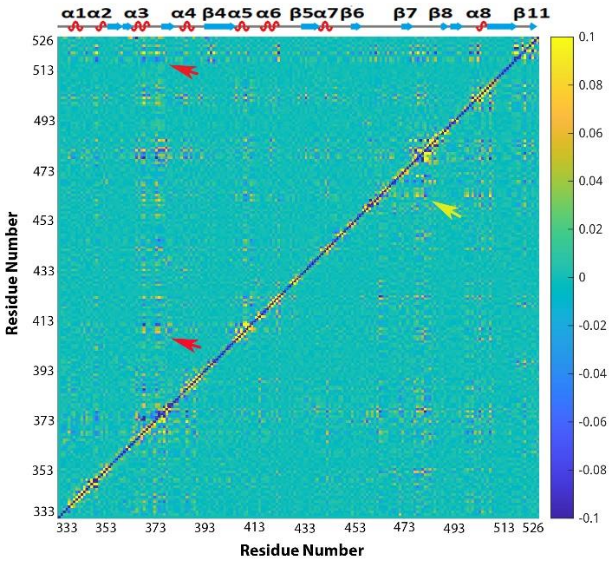


Figure S10. Comparison of the averaged mixed dihedral angle covariance matrix for RBD-K417N (upper-left triangle above the diagonal line) in comparison of RBD in wildtype (lower-right triangle) in bound state (Left). The result from RBD in wildtype is shown in the lower-right triangle below the diagonal line.

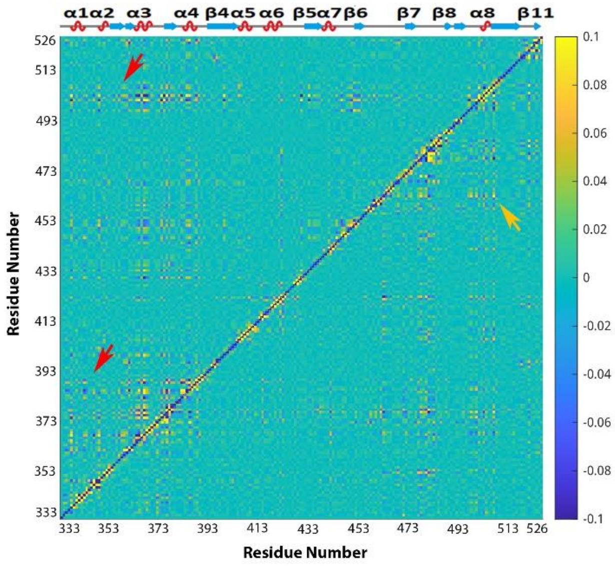


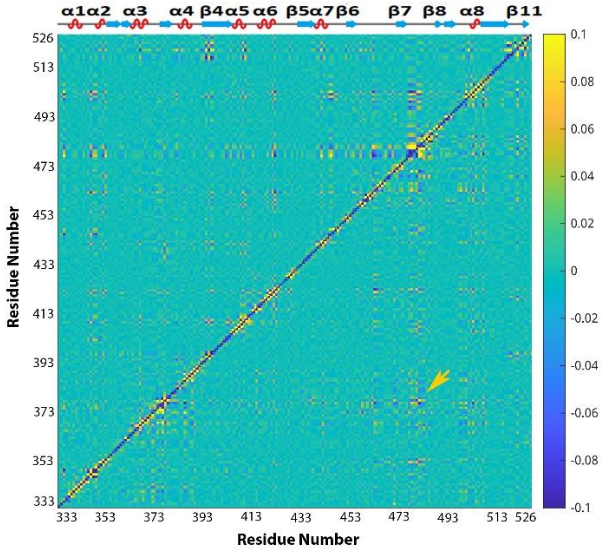


Figure S11. Comparison of the averaged mixed dihedral angle covariance matrix for RBD-N439K (upper-left triangle above the diagonal line) in comparison of RBD in wildtype (lower-right triangle) in bound state (Left), and RBD-P479S (upper-left triangle above the diagonal line) in comparison of RBD in wildtype (lower-right triangle). The result from RBD in wildtype is shown in the lower-right triangle below the diagonal line.

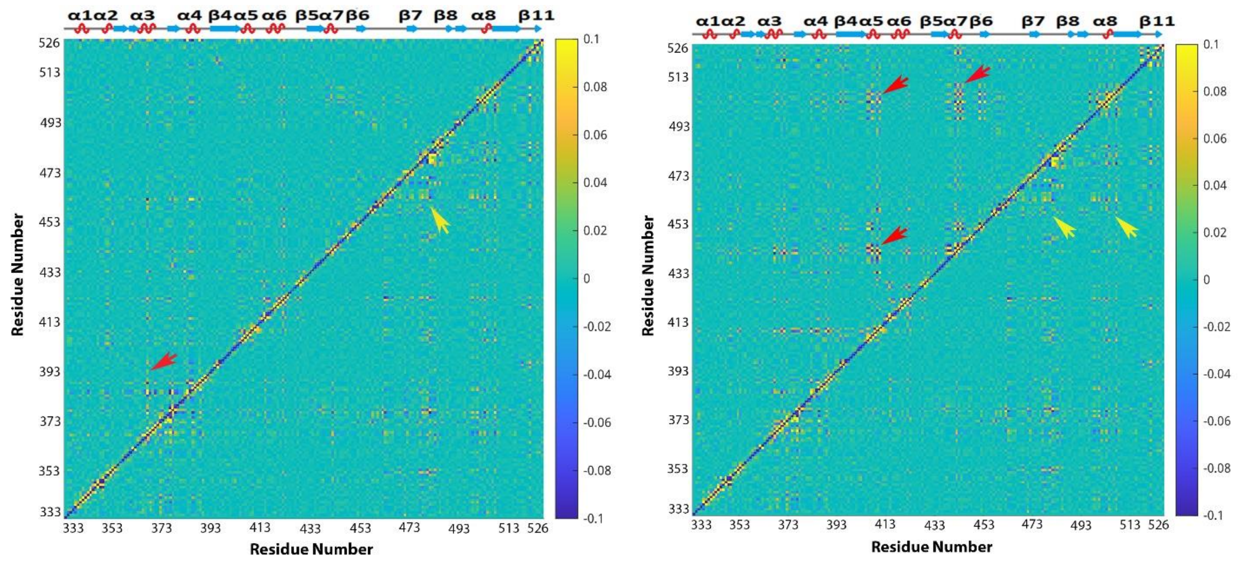


Figure S12. Comparison of the averaged mixed dihedral angle covariance matrix for RBD-S477N (upper-left triangle above the diagonal line) in comparison of RBD in wildtype (lower-right triangle) in bound state (Left), and RBD-T478I (upper-left triangle above the diagonal line) in comparison of RBD in wildtype (lower-right triangle). The result from RBD in wildtype is shown in the lower-right triangle below the diagonal line.

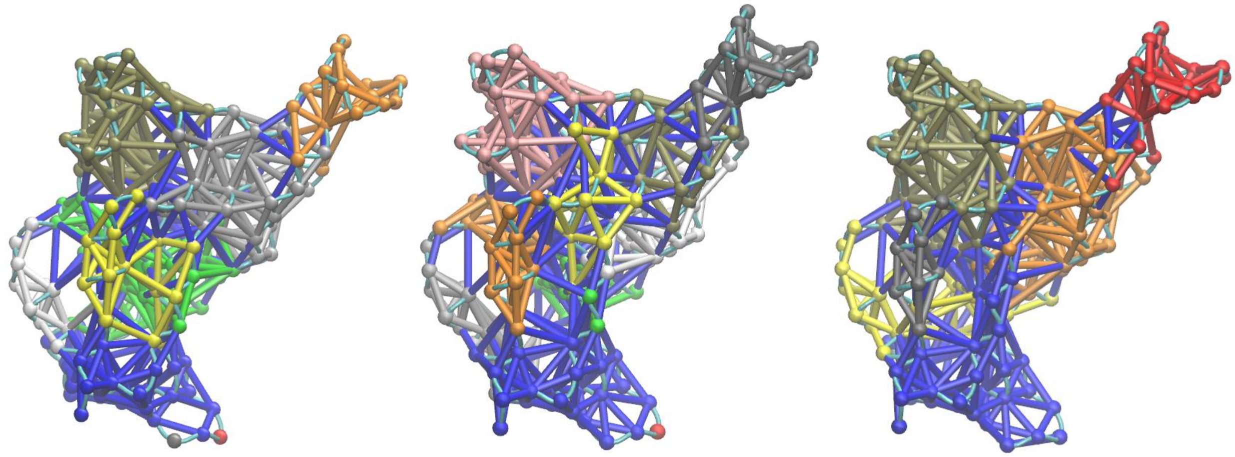

Figure S13. Community networks formed in the RBD-E484K(Left), in RBD-K417N (middle) and RBD-N439K (Right) based on MD simulation and dynamical network analysis. The RBD network structures are oriented in the same direction as the RBD shown in Figure 1.

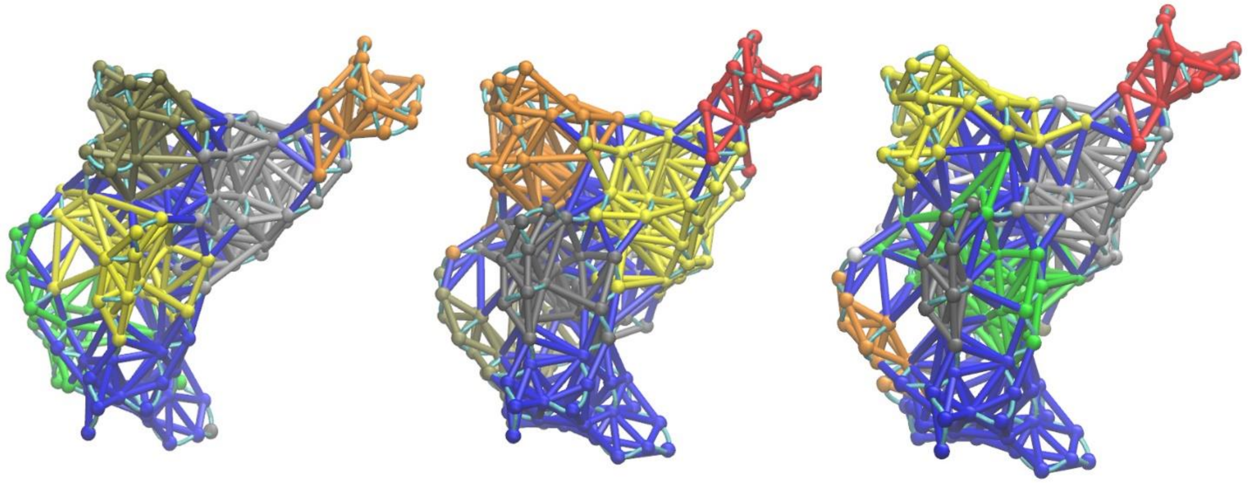

Figure S14. Community networks formed in the RBD-P479S(Left), in RBD-S477N (middle) and RBD-T478I (Right) based on MD simulation and dynamical network analysis. The RBD network structures are oriented in the same direction as the RBD shown in Figure 1.
